# Amphicarpic development in the emerging model organism *Cardamine chenopodiifolia*

**DOI:** 10.1101/2024.01.26.577352

**Authors:** Aurélia Emonet, Miguel Pérez-Antón, Ulla Neumann, Sonja Dunemann, Bruno Huettel, Robert Koller, Angela Hay

## Abstract

- Amphicarpy is an unusual trait where two fruit types develop: one above and the other below ground. This trait is not found in conventional model species, therefore, its development and molecular genetics remain under-studied. Here, we establish *Cardamine chenopodiifolia* as an emerging experimental system to study amphicarpy.
- We characterized the development of *C. chenopodiifolia*, focusing on differences in morphology and cell wall histochemistry between above- and below-ground fruit. We generated a reference transcriptome using PacBio full-length transcript sequencing (IsoSeq) and used a combination of short and long read sequencing to analyse differential gene expression between above- and below-ground fruit valves.
- *C. chenopodiifolia* has two contrasting modes of seed dispersal. The main shoot fails to bolt and initiates floral primordia that bury underground where they self-pollinate and set seed. By contrast, axillary shoots bolt to position flowers and exploding seed pods above ground. Morphological differences between aerial explosive fruit and subterranean non-explosive fruit were reflected in a large number of differentially regulated genes involved in photosynthesis, secondary cell wall formation and defence responses.
- Tools established in *C. chenopodiifolia*, such as a reference transcriptome, draft genome assembly and stable plant transformation, pave the way to explore under-studied traits and discover new biological mechanisms.

## Introduction

Seed dispersal strategies vary widely between different plants. In flowering plants, seeds are encased in fruit for both protection and dispersal, and fruit diversity reflects the many different adaptations for dispersal seen in nature. Fruit of the model plant Arabidopsis are dry siliques that open by dehiscence and seeds are released once the fruit structure falls apart (Dinneny *et al*., 2005). Genetic tools in Arabidopsis have been immensely useful for studying fruit development and the tissue patterning required for dehiscence (Liljegren *et al*., 2000, 2004; Ferrándiz *et al*., 2000). However, only a fraction of the diverse seed dispersal strategies found in nature are accessible through the study of model organisms.

Comparative approaches extend the reach of developmental genetics by developing new tools in less well-studied species. For example, *Cardamine hirsuta* is a relative of Arabidopsis with many divergent traits and genetic tools to study them (Hay & Tsiantis, 2006; Hay *et al*., 2014; Vlad *et al*., 2014; Monniaux *et al*., 2018; Baumgarten *et al*., 2023). Explosive seed dispersal is one such trait that distinguishes *C. hirsuta* from Arabidopsis. Seeds are launched at speeds greater than 10 ms^-1^ by the explosive coiling of *C. hirsuta* fruit valves (Hofhuis *et al*., 2016). This rapid movement disperses seeds across a large area, which is an efficient strategy for a ruderal weed.

Explosive seed dispersal is a derived trait, shared by *Cardamine* species in the Brassicaceae. A novel pattern of lignified secondary cell walls evolved in strict association with this trait (Hofhuis *et al*., 2016). A hardened inner layer called the endocarp typically forms in the valves of all Brassicaceae fruit. These lignified cells form the endocarp *b* cell layer in Arabidopsis and *C. hirsuta*. These cells are uniformly lignified in Arabidopsis and other species with non-explosive fruit (Spence *et al*., 1996). In contrast to this, a thick, lignified wall is deposited asymmetrically in a distinctive hinged pattern in the endocarp *b* cells of *Cardamine* species with explosive fruit (Hofhuis *et al*., 2016). Genetics and computational modelling approaches in *C. hirsuta* have shown that this asymmetric pattern of lignin deposition provides a mechanism to rapidly release built-up tension in the valves via explosive coiling (Hofhuis *et al*., 2016; Pérez Antón *et al*., 2022).

Seed dispersal strategies can also vary within a single plant through the development of two or more fruit/seed morphs. This heteromorphy is exhibited by hundreds of flowering plant species and often described as a bet-hedging strategy (Imbert, 2002; Gianella *et al*., 2021). Plants ‘hedge their bets’ by following more than one dispersal strategy to increase their chances of success in variable environments. In the Brassicaceae, for example, *Diptychocarpus strictus* and *Aethionema arabicum* develop both dehiscent and indehiscent fruit on the same plant (Lu *et al*., 2010, 2015; Lenser *et al*., 2016). The dehiscent fruit in *Aethionema arabicum* release mucilaginous seeds that germinate quickly, while indehiscent fruit produce a single seed that is not released and has delayed germination (Lenser *et al*., 2016).

Amphicarpy is a specific type of heteromorphy where one fruit morph is buried underground. Underground fruit, or geocarpy, is a strategy used by a number of species for seed dispersal, such as the agronomically important peanut plant (*Arachis hypogaea,* Chen et al., 2016). In contrast to this, amphicarpy is a dual strategy, combining the advantages of both aerial and subterranean fruit for seed dispersal (Cheplick, 1987). Amphicarpy is a rare trait that evolved independently in different groups of flowering plants, particularly in the Fabaceae family (Kaul *et al*., 2000; Zhang *et al*., 2020a; Liu *et al*., 2021). For example, the legume *Amphicarpaea edgeworthii* is an amphicarpic plant with a recently sequenced genome (Liu *et al*., 2021). The Brassicaceae family contains one amphicarpic species that belongs to the *Cardamine* genus. *Cardamine chenopodiifolia* Pers. is an annual plant, native to South America, that is easily cultivated in greenhouse conditions (Persoon, 1807; Gorczyński, 1930; Cabrera, 1967; Cheplick, 1983). A recent draft assembly of the octoploid genome of *C. chenopodiifolia* provides some of the necessary tools to develop this plant as an emerging experimental system to study amphicarpy (Emonet *et al*., 2024).

An interesting feature of *C. chenopodiifolia* is that it combines both explosive and non-explosive seed dispersal. It partitions these two very different modes of seed dispersal between explosive aerial fruit and non-explosive subterranean fruit. Comparing both traits within the same plant provides a complementary approach to comparisons between Arabidopsis and *C. hirsuta*. In particular, comparative transcriptomics between two fruit morphs encoded by a single genome, bypasses many of the challenges inherent to between-species comparisons. For these reasons, we characterised amphicarpic development in *C. chenopodiifolia,* focusing on the differences in morphology, cell wall histochemistry and gene expression profiles that differentiate aerial and subterranean fruit. This work establishes resources in *C. chenopodiifolia* as an emerging experimental system, and identifies differences in secondary wall deposition in the fruit endocarp between the two fruit types as important determinants of explosive versus non-explosive seed dispersal.

## Materials and Methods

### Plant and growth conditions

*Cardamine chenopodiifolia* Pers. seeds (Ipen: XX-0-MJG-19—35600) were obtained from the Botanic Garden of the Johannes Gutenberg University, Mainz, Germany. To improve germination, aerial seeds were germinated in long-days on ½ Murashige and Skoog (MS) plates after 7 days stratification, then 1-week-old seedlings were transferred to soil and grown in long-day (16h light, 20 °C; 8h dark, 18°C; 65% humidity) or short-day conditions (8h light, 20 °C; 16h dark, 18°C, 65% humidity). See Methods S1 for details. For Magnetic Resonance Imaging (MRI), seedlings were transferred to Speyer 2.1 soil (loamy sand, sieved to 2 mm, brand name Sp2.1, Landwirtschaftliche Untersuchungs-und Forschungsanstalt Speyer, Speyer, Germany) and grown in a greenhouse under long-day conditions.

### Plant transformation

*DR5v2::NLS-tdTomato* construct in a modified pPZP200 vector (Bhatia *et al*., 2023) was transformed into *C. chenopodiifolia* by floral dip (Clough & Bent, 1998). Eighty plants were transformed and T1 transformants were selected with basta treatment (Methods S1).

### Seed, fruit and plant phenotyping and staging

Forty plants were grown over a period of 6 weeks and their characteristics monitored every one or two days (see Methods S1 for details). At 44-53 days post-germination (dpg), 17 plants were used to count aerial and subterranean fruit, measure fruit width and length, and ascribe fruit stages. A MARVIN seed counter was used to count and measure seed weight and dimensions of dry aerial and subterranean seeds from 4 plants (Methods S1).

### Heteroblastic leaf shape analysis

Leaves of 7-week-old plants (n=4) were flattened onto white paper using transparent adhesive film and scanned with an Epson Perfection V700 Photo scanner. Morphometric analysis was carried out on leaf silhouettes using the LeafI software (Leaf Interrogator, Z. Zhang et al., 2020).

### Mucilage assay

Seed mucilage was stained with 0.01% ruthenium red solution (McFarlane *et al*., 2014) and imaged under a Niko SMZ18 binocular with SHR Plan Apo 1.6x WD30 objective.

### Photography

Photographs of plants at different stages were taken with a Nikon D800 equipped with either AF-S Micro NIKKOR 105mm 1:2.8G ED or AF-S NIKKOR 24-85mm 1:3.5-4.5 G objectives.

### Microscopy

Brightfield microscopy was performed using a Zeiss Axio Imager M2 upright microscope with EC Plan-Neofluar 10x/0.3, 20x/0.5 or 40x/0.75 objectives. Confocal laser scanning microscopy (CLSM) was performed on a Leica TCS SP8 with HC PL FLUOTAR (10x/0.30 dry) and HCX PL APO lambda blue (63x/1.20 water) objectives. Excitation and detection windows were set as follows: for visualization of lignin and cellulose: calcofluor (405nm, 425-475nm), basic fuchsin (561nm, 600-650nm). Images were processed using Fiji software (Schindelin *et al*., 2012). Scanning electronic microscopy was performed using a Zeiss Supra 40VP microscope.

### Histochemistry

For toluidine blue staining, samples were fixed, embedded in LR White, sectioned, stained and imaged by light microscopy (Neumann & Hay, 2019). To visualize lignin by CLSM, 100-150um transverse sections were cut using a Leica Vibratome VT1000 S, immediately fixed and cleared using an adapted Clearsee protocol with Basic Fuchsin and Calcofluor white staining (Ursache *et al*., 2018; Pérez Antón *et al*., 2022). For SEM, plant material was fixed in 4% glutaraldehyde, dehydrated through an increasing ethanol series and critical point dried using a Leica CPD300 critical point dryer. Samples were mounted and sputter coated with platinum using a Polaron SC 7640 sputter coater (Methods S1).

### High speed video

Explosive pod shatter was filmed with a high-speed camera (Photron Fastcam SA3 120KM2, Photron Europe Ltd, Bucks, UK) fitted with a 100 mm F2.8 lens and recorded at 10000 frames per second (Methods S1).

### Time lapse photography of subterranean fruit

Four-week-old *C. chenopodiifolia* plants were grown in a transparent chamber and roots and subterranean fruit were imaged every 30 min with a Canon EOS 700D camera equipped with an EF-S 18-55mm f/3.5-5.6 lens (Methods S1).

### Magnetic resonance imaging (MRI)

MRI was performed at Jülich Forschungszentrum GmbH, Jülich, Germany using a vertical bore 4.7T magnet (Magnex UK), on ten-week-old plants grown in soil whose humidity was maintained constant at 70% of maximum water holding capacity (WHCmax) (Pflugfelder *et al*., 2022). Image resolution, 0.3×0.3×0.6 mm^3^, Echo time, 9 ms, repetition time 2s, field of view 77×77×84 mm^3^.

### RNA Sequencing

RNA of aerial and subterranean fruit valves (stage 16-17a and 17ab) was extracted from three biological replicates using Spectrum^TM^ Plant Total RNA kit (Sigma-Aldrich). For long read sequencing, full-length complementary DNA (cDNA) synthesis was performed with TeloPrime Full-Length cDNA Amplification Kit (Lexogen) (Cartolano *et al*., 2016). A Single Molecule Real-Time (SMRT) bell library (Pacific Biosciences) was prepared with the IsoSeq protocol and sequenced on PacBio Sequel IIe at the Max Planck Institute for Plant Breeding Research Genome Centre. PacBio IsoSeq data were demultiplexed and used to build and annotate a *C. chenopodiifolia* fruit reference transcriptome using SMRTLink (v10.2.0.133434), TAMA (tc_version_date_2019_11_19) and *C. chenopodiifolia* genome assembly information (Kuo *et al*., 2020; Emonet *et al*., 2024).

For short read sequencing, the cDNA synthesis, Stranded Poly-A selection library preparation (two-sided, 150bp) and sequencing were carried out by Novogene using the Illumina NovaSeq6000 platform. Paired-end short reads were quality-checked, aligned to both the *C. chenopodiifolia* fruit transcriptome and *C hirsuta* genome (Gan *et al*., 2016) and quantified using Salmon (v1.8.0) and HISAT2 (v2.1.0), respectively (Patro *et al*., 2017; Kim *et al*., 2019). Differential expression analysis was performed with EdgeR (v3.42.4) and DESeq2 (v1.36.0) (Love *et al*., 2014; Chen *et al*., 2016b), respectively. Cluster and Gene Ontology (GO) analyses were carried out using ClusterProfiler (v.4.4.4) (Wu *et al*., 2021). GO analysis was performed using *A. thaliana* GO term annotations for orthologous *C. hirsuta* genes (Gan *et al*., 2016). For more details see Methods S1.

### Statistical analysis

All statistical analyses were carried out using R (v4.2.1) and RStudio (v. 2022.07.1) (‘R’, 2022) (Methods S1).

### Data Availability

Short-sequence read data and PacBio long-sequence read data for this study has been deposited in the European Nucleotide Archive (ENA) at the European Molecular Biology Laboratory’s European

Bioinformatics Institute (EMBL-EBI) under accession number PRJEB69676 (https://www.ebi.ac.uk/ena/browser/view/PRJEB69676)

## Results

### Amphicarpy: above- and below-ground fruit

Two different fruit morphs develop in *C. chenopodiifolia*: one type above ground and the other below ground (Fig. 1). Subterranean fruit are derived from flowers that develop from the shoot apical meristem (Fig. 1B-D). Floral buds are produced at the shoot apical meristem after three to four weeks of growth in long-day conditions (Fig1B, Fig S1B). Each floral bud is immediately propelled into the soil by differential growth and rapid elongation of its pedicel (Fig 1B-D, Movie 1). Gravitropic growth of the pedicel continues, reaching an average length of 5 cm (Fig. 1E, G). About twenty subterranean fruits develop in our standard growth conditions (Fig. 1F). The first fruit produced usually grow close to the main root, while later fruit spread out around the periphery of the root system (imaged *in situ* by MRI, Movie 2). At this stage of development, the shoot apex looks like an octopus when viewed from above, with pedicels extending down and anchoring the rosette plant into the soil (Fig. 1C). These pedicels have large cortical cells with Casparian strip-like lignin deposition in the innermost cortical cell layer surrounding the vasculature, which is absent in the pedicels of aerial fruit (Fig.1H, S1E). Aerial fruit, on the other hand, are derived from flowers that develop from axillary meristems (Fig. 1A-B). Soon after the transition to flowering, the growth of meristems already formed in the axils of rosette leaves is released (Fig. 1B, S1A-B). Five to six of these axillary shoots grow out and aerial flowers are first observed at 45 days post germination (Fig. S1A-B). An average of 24 aerial fruits develop from the flowers on each shoot, excluding additional fruit that form on secondary branches (Fig. 1F).

**Fig. 1.**
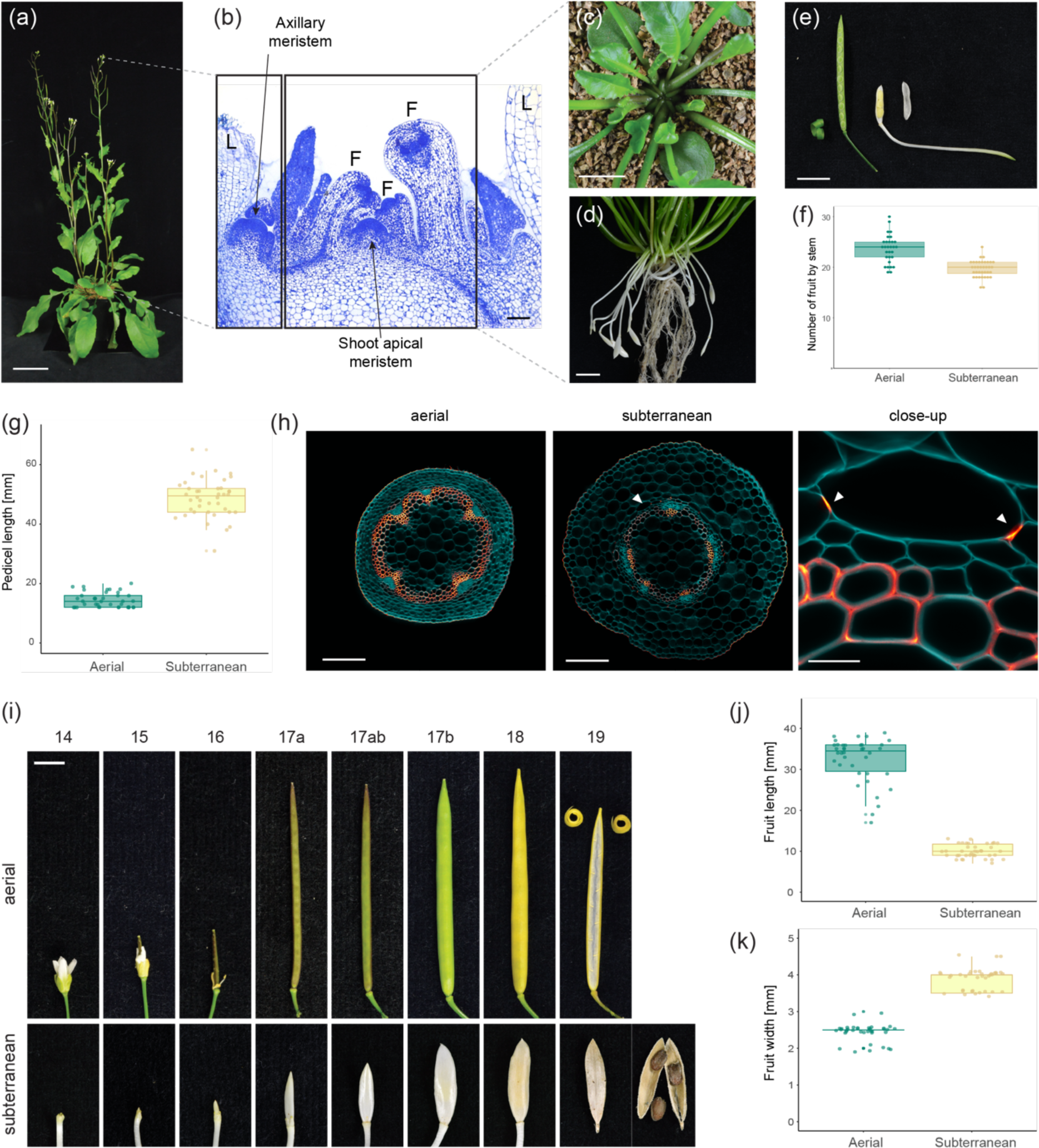
*Cardamine chenopodiifolia* is amphicarpic. (a) *C. chenopodiifolia* plant at 60 dpg. (b) Toluidine blue-stained longitudinal section showing shoot apical meristem and axillary meristem. F, flower; L, leaves. (c) Top view of shoot apex of 4-week-old plant, with flower pedicels growing into the soil and axillary shoots growing out of the rosette. (d) Subterranean flowers and fruit borne on long pedicels, and roots, after soil removal from a 7-week-old plant. (e) Aerial (left) and subterranean (right) fruit and pedicels with one valve removed. Aerial fruit valve coils, while subterranean valve does not. (f-g) Box plots of fruit number per stem (f) and pedicel length (g) in aerial and subterranean fruits. Plots show median (thick horizontal lines), n = 30-40 plants at approximately 80 dpg (dots). Differences between fruit morphs were analysed using Wilcoxon test, p-val = 1.111e-07 (f) and p-val = 2.711e-14 (g). (h) Confocal laser scanning micrographs (CLSM) of 100 µm transverse sections of aerial and subterranean pedicels stained with calcofluor white (cellulose, cyan) and basic fuchsin (lignin, Red Hot LUT). Localized lignin deposition in the innermost cortical cell layer surrounding the vasculature in subterranean pedicels, indicated by white arrowheads and in close-up view. (i) Aerial and subterranean flowers and fruit at stage 14 through 19. (j-k) Box plots of fruit length (j) and width (k) in aerial and subterranean fruits. Plots show median (thick horizontal lines), n = 38-40 fruit per morph (dots). Differences between fruit morphs were analysed using Wilcoxon test, p-val = 5.558e-14 (j) and p-val = 4.214e-15 (k). Bars: 10 cm (a), 100 µm (b), 1 cm (c, d, e), 200 µm (h), 20 µm (close-up, h), 5 mm (i).

Both of the fruit morphs in *C. chenopodiifolia* are dehiscent siliques, but they employ different mechanisms of seed dispersal (Fig. 1E). Seeds are dispersed over a large area by explosive coiling of the two valves in aerial fruit (Movie 3). The first aerial fruit explode after 11 weeks of growth in long day conditions (Fig. S1B). By contrast, subterranean fruit develop over a much longer period and dehisce non-explosively to release their seeds underground long after all aerial fruit have exploded.

The two fruit morphs look very different: aerial fruit are elongated and green, while subterranean fruit are short and white (Fig. 1E, J-K). The development of both fruit morphs follows a similar progression, elongating and then expanding in girth before the fruit tissues stiffen due to lignin deposition at stage 17b (Fig. 1I). However, the final stages of development are very different between the two morphs. Aerial fruit explosively release their seeds after stage 17b, while subterranean fruit continue to develop through to stage 19, at which point they dry and dehisce (Fig. 1I).

### Endocarp *b* lignification differs between fruit morphs

Explosive coiling of the aerial fruit valves is associated with a specific pattern of secondary cell wall lignification (Fig. 2). The valves of both fruit morphs comprise one exocarp cell layer, several layers of mesocarp cells and two endocarp cell layers, *a* and *b* (Fig. 2A-B). Endocarp *b* cells have a lignified secondary cell wall and the patterning of this wall differs dramatically between the two fruit morphs. In aerial fruit, a thick secondary cell wall, disrupted by two thin hinges, is deposited asymmetrically in endocarp *b* cells (Fig. 2A-B). By contrast, subterranean fruit have 1 to 3 endocarp *b* cell layers in which a thick secondary cell wall is deposited uniformly (Fig. 2A-B).

**Fig. 2.**
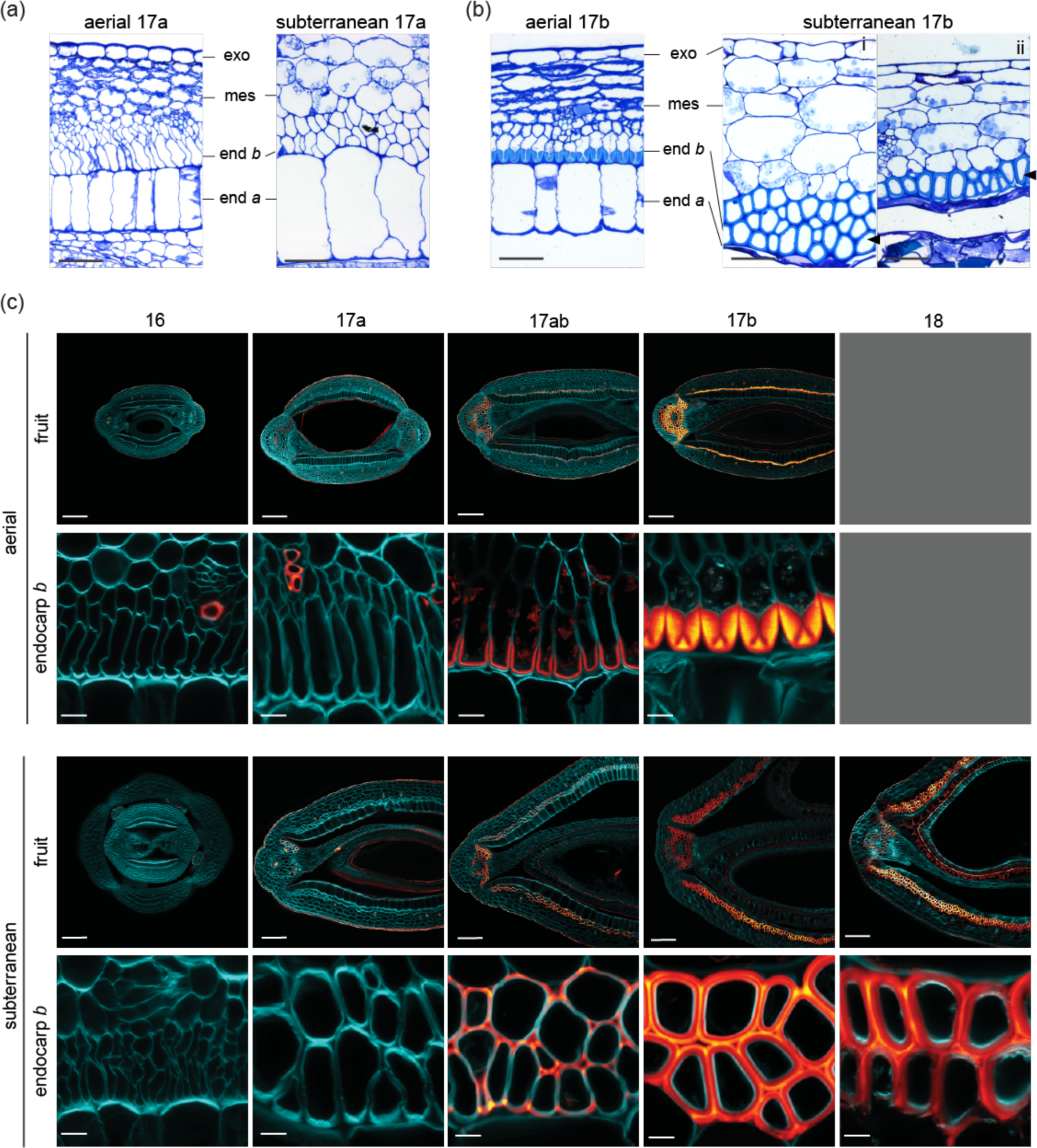
Secondary cell wall patterning in endocarp *b* cells differs between aerial and subterranean fruit. (a-b) Toluidine blue-stained transverse sections of aerial and subterranean fruit valves at stage 17a (a, before lignification) and 17b (b, after lignification). Exo, exocarp; mes, mesocarp; end *b*, endocarp *b*, end *a*, endocarp *a*. Arrows indicate endocarp *b* (b). (c) CLSM of 100 µm transverse sections of aerial and subterranean fruit at stage 16 through 18 stained for cellulose (calcofluor white, cyan) and lignin (basic fuchsin, Red Hot LUT). Aerial fruit could not be sectioned at stage 18 due to explosive coiling. Bars: 50 µm (a, b), 100 µm (c, fruit), 10 µm (c, endocarp *b*).

Starting from a similar fruit structure at stage 16, the pattern of lignin deposition diverges between aerial and subterranean fruit valves after stage 17a (Fig. 2C). In aerial fruit, endocarp *b* cells form a lignified secondary cell wall solely on the adaxial side (Fig. 2C). In these cells, lignin is deposited in a thin line that forms a “U” in cross section at stage 17ab (Fig. 2C). By stage 17b, this lignified wall is very thick, but disrupted by two thin hinges at the base of the “U” (Fig. 2C).

In subterranean fruit, the uniform lignification of endocarp *b* cells initiates in cell corners at stage 17ab, but does not fill the three-way junctions between cells (Fig. 2C). Lignin deposition then continues throughout the cell wall, resulting in a uniformly thickened secondary cell wall by stage 17b (Fig. 2C). Notably, these stages of fruit development are considerably longer in subterranean than aerial fruit. Multiple endocarp *b* cell layers can form in subterranean fruit valves, and although these cell layers are present in aerial fruit valves, they do not differentiate a lignified secondary cell wall (Fig. 2A-B). Therefore, the two fruit morphs differ not only in the initiation and patterning of endocarp *b* lignification, but also in the specification of endocarp *b* cell fate.

As subterranean fruit desiccate from stage 18 onwards, a small amount of lignin is deposited in mesocarp cell walls that are adjacent to endocarp *b* cells (Fig. 2C). Lignification of valve margin cells also begins at this stage in order to form a dehiscence zone between the replum and each valve (Fig. 2B, S2A-B). In aerial fruit, by contrast, valve margin cells are fully lignified much earlier at stage 17b, when the lignification of endocarp *b* and replum cells is also completed (Fig. 2B, S2A-B). Another conspicuous difference between the two fruit morphs is the sudden collapse of endocarp *a* cells between stages 17a and 17b in the valves of subterranean fruit, but not aerial fruit (Fig. 2A-B).

### Seeds and flowers differ between fruit morphs

Seeds that develop in aerial versus subterranean fruit are distinguished by many characteristics related to their different modes of dispersal. Each aerial fruit contains around 20 seeds compared to an average of only 2 seeds in subterranean fruit (Fig. 1D, Fig. 3B). These subterranean seeds are larger and heavier than aerial seeds (Fig. 3B-E). Aerial seeds are shaped like flattened discs compared to the oval shape of subterranean seeds (Fig. 3B, F). Moreover, aerial seeds release a large amount of mucilage, compared to subterranean seeds, upon imbibition (Fig. 3G). This is reflected in a large difference in the amount of mucilage produced in epidermal cells of the seed coat in aerial compared to subterranean seeds (Fig. 3H). Other features, such as the seed coat surface and the seed abscission zone appear very similar between both seed types (Fig. S2C-E). Therefore, aerial fruit produce numerous, small, mucilaginous seeds, suitable for long range dispersal. In contrast, subterranean fruit produce few, large seeds, which is thought to aid germination from greater soil depths (Cheplick, 1987; Zhang *et al*., 2020a).

**Fig. 3.**
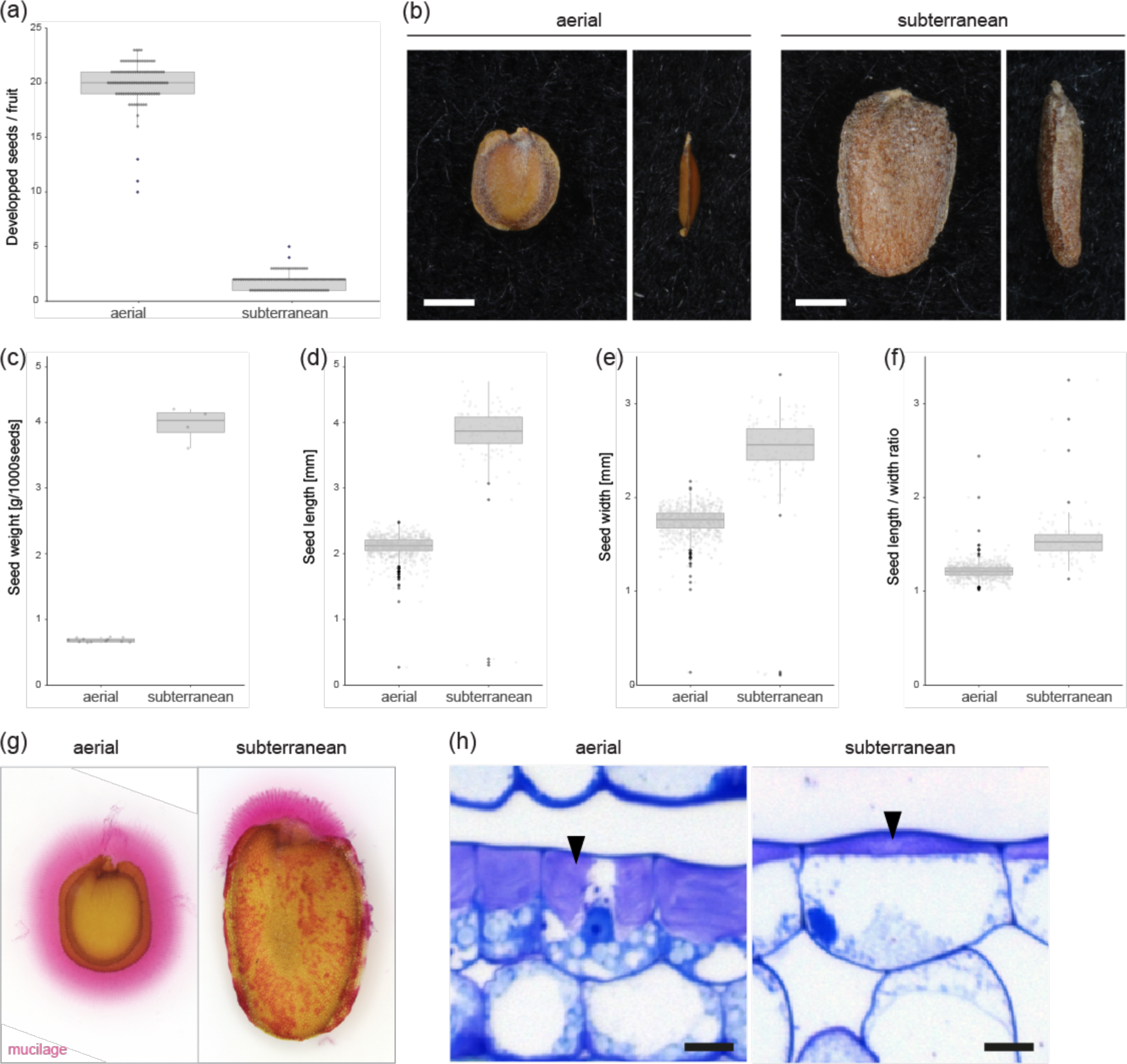
Differences between aerial and subterranean seeds. (a) Box plot of seed number per fruit in aerial and subterranean fruits. Plot shows median (thick horizontal lines), n = 100 seeds per morph (dots). Differences between fruit morphs were analysed using Wilcoxon test, p-val = 2e-16. (b) Aerial and subterranean seeds shown from front (left) and side (right) views. (c-f) Box plots of seed weight per 1000 seeds (c), seed length (d), seed width (e) and seed length/width (f) in aerial and subterranean fruits. Plots show median (thick horizontal lines), n = 115 subterranean and 1970 aerial seeds from 4 plants (dots). Differences between seed morphs were analysed using Wilcoxon test, p-val = 0.0011 (c), p-val = <2e-16 (d), p-val = <2e-16 (e), p-val = <2e-16 (f). (g) Ruthenium red-stained mucilage (pink) released by aerial and subterranean seeds after imbibition. (h) Toluidine blue-stained transverse sections of outer seed coat of aerial and subterranean seeds from stage 17b fruit. Mucilage pockets indicated by black arrows. Bars: 1 mm (b, g), 10 µm (h).

Both flower morphs in *C. chenopodiifolia* are capable of selfing, but subterranean flowers employ an automatic type of self-pollination called cleistogamy. Aerial flowers are typical of the Brassicaceae, with a perianth of 4 green sepals and 4 white petals, enclosing 6 stamens and two carpels (Fig. 4A-B). Subterranean flowers, on the other hand, are non-opening (Fig. 4A). The 4 sepals remain closed, protecting the developing reproductive organs as the flower pushes through the soil (Movie 1). Removing the sepals in stage 14 flowers reveals the pollen-bearing anthers in direct contact with the stigmatic papillae of the gynecium (Fig. 4C). In addition, the floral organs of subterranean flowers are very reduced in size compared to aerial flowers (Fig. 4A, C) and fewer than 4 petals are sometimes observed (Fig. 4C, S2F).

**Fig. 4.**
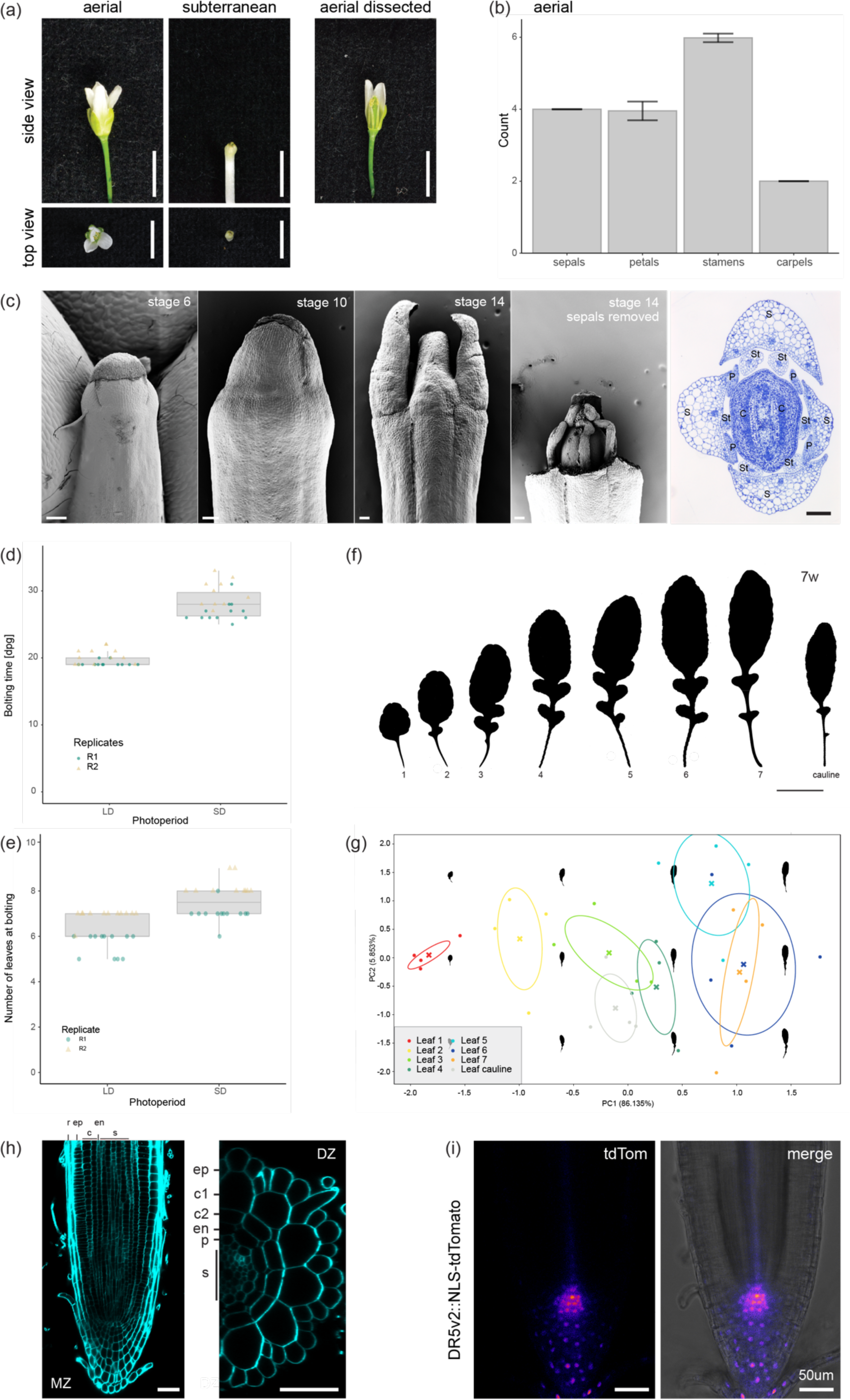
Above- and below-ground organs in *C. chenopodiifolia*. (a) Aerial and subterranean flowers at stage 14, shown in side and top views. Perianth organs dissected off an aerial flower to show reproductive organs. (b) Bar plot of the number of sepals, petals, stamens and carpels in aerial flowers; plot shows mean and standard deviation, n = 87. (c) Scanning electron micrographs of subterranean flowers at stages 6, 10 and 14; sepals dissected off a stage 14 flower to show reproductive organs. Toluidine blue-stained transverse section of subterranean flowers at stage 14; S, sepals; P, petals; St, stamens; C, carpels. (d-e) Box plots of bolting time in day post germination (dpg) (d), and number of leaves at bolting time (e) for plants grown in long day (LD) or short day (SD) conditions. Plots show median (thick horizontal lines), n = 24 plants per photoperiod (dots). Differences between photoperiods were analysed using Wilcoxon test, p-val = 3.811e-09 (d), p-val = 6.579e-06 (e). (f) Heteroblastic leaf series of 7-week plant. (g) Morphometric analysis of heteroblastic leaf series of 7-week plant (n = 4 plants). (h) CLSMs of seedling root tip meristematic zone (MZ, left) and optical section of root differentiation zone (DZ, right); r, root cap; ep, epidermis; c, cortex; c1, first cortex cell layer; c2, second cortex cell layer; en, endodermis; p, pericycle; s, stele. (i) CLSM of *DR5v2::NLS-tdTomato* expression (Fire LUT, left) and merged with brightfield image (right) in seedling root tip. Bars: 5 mm (a), 100 µm (c), 5 cm (f), 50 µm (h, i).

### Plant architecture above and below ground

*Cardamine chenopodiifolia* is a long-day plant. By measuring time to flowering and the number of rosette leaves at flowering, we found that flowering was accelerated in long-day compared to short-day photoperiods (Fig. 4D-E). While the axillary shoots of *C. chenopodiifolia* produce long stems that position flowers far above the rosette, flowers that are produced at the shoot apical meristem are positioned underground by elongating pedicels that grow towards gravity (Fig. 1A-C, Fig. S1A-B). The main shoot does not bolt and has a rosette flowering habit, while axillary shoots produce elongated internodes, cauline leaves and secondary branches (Fig. 1A, Fig. S1A, C).

The rosette of *C. chenopodiifolia* generally comprises 7 large, lobed leaves, arranged in a spiral phyllotaxy (Figs. 1A, 4F, Fig. S1C). Heteroblasty is not particularly pronounced, but morphometric analysis clearly separates the rounder shape of the first leaves from the increasingly more oval and lobed shape of later leaves (Fig. 4F-G).

Roots of *C. chenopodiifolia* have a similar tissue structure to *C. hirsuta* (Hay *et al*., 2014), comprising an epidermis, two cortex layers, an endodermis, a pericycle and vascular tissues filling the central cylinder (Fig. 4H). The quiescent centre is marked by a maximum of auxin activity, as reported by *DR5v2::NLS-tdTomato* (Fig. 4I). This transgene was stably transformed by floral dip of aerial flowers using *Agrobacterium tumefaciens* (see Materials and Methods), demonstrating that transgenic approaches can be followed in *C. chenopodiifolia*.

In summary, *C. chenopodiifolia* is an amphicarpic plant that uses two very distinct strategies for seed dispersal. These different strategies are associated with a suite of distinct traits that differentiate the development of reproductive structures above and below ground. For these reasons, *C. chenopodiifolia* provides a unique framework to compare morphological divergence between two fruit morphs that develop on the same plant.

### Comparative transcriptome analysis

To compare how the transcriptome of *C. chenopodiifolia* is differentially regulated in aerial versus subterranean fruit valves, we performed RNA-seq using both long read and short read sequencing. First, we used Pacific Bioscience (PacBio) Single-Molecule Real-Time (SMRT) long read isoform sequencing (IsoSeq) to generate a reference transcriptome for *C. chenopodiifolia* fruit valves (Fig. S3A). We sequenced RNA of aerial and subterranean fruit valves harvested at two developmental stages (late stage 16 and stage 17ab), during which the valves reach their final length. These stages correspond to before and during the development of trait differences between the fruit types, such as endocarp *b* lignin patterning and endocarp *a* collapse (Fig. 2). The four sample types were collected from the same plant, valves from five different plants were pooled to ensure sufficient material, and the experiment was performed in triplicate (Fig. 5A). To increase gene diversity in our reference transcriptome, we sequenced additional samples harvested from different tissue types and hormone/elicitor treatments (Methods S1). PacBio IsoSeq provided high quality isoforms (Table S1) that we collapsed into a reference transcriptome using information from a draft genome assembly of *C. chenopodiifolia* (Table S1) (Emonet *et al*., 2024). We annotated 53242 genes in this reference transcriptome using the Uniref90 protein database. Approximately 94% of full-length, non-concatemer reads (FLNC) aligned to the draft *C. chenopodiifolia* genome (Emonet *et al*., 2024), suggesting that almost all of our reference transcriptome is included in this draft genome assembly.

**Fig. 5.**
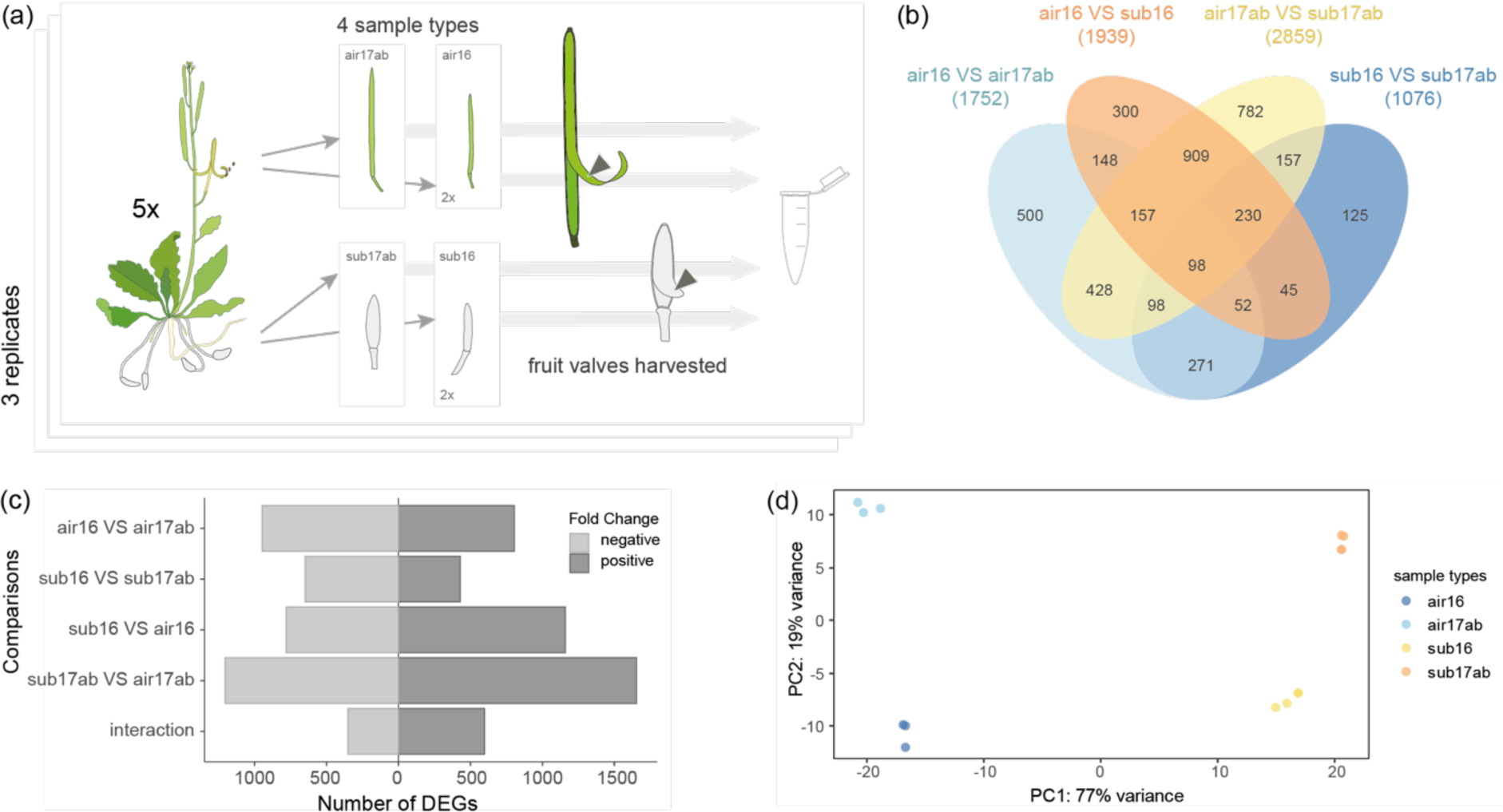
Transcriptomes differ markedly between aerial and subterranean fruit valves. (a) Schematic of RNA-seq design. (b) Venn diagram of differentially expressed genes (DEG, logFC ≥│1│, padj < 0.05) between aerial fruit stage 16 VS aerial fruit stage 17ab (air16 VS air17ab), aerial fruit stage 16 VS subterranean fruit stage 16 (air16 VS sub16), aerial fruit stage 17ab VS subterranean fruit stage 17ab (air17ab VS sub17ab), subterranean fruit stage 16 VS subterranean fruit stage 17ab (sub16 VS sub17ab). DEGs were obtained by mapping *C. chenopodiifolia* short reads to the *C. hirsuta* genome. (c) Number of up- and down-regulated genes for each comparison (DEG, logFC ≥│1│, padj < 0.05). (d) Principal component analysis of short read data from *C. chenopodiifolia* fruit valve sample types.

To identify genes that are differentially expressed between aerial and subterranean fruit valves, we used the same twelve RNA samples from *C. chenopodiifolia* valves for short read sequencing. Illumina sequencing provided 89.7Gb of raw data with an average of 50 million reads per sample (Table S2). We aligned these short reads to the *C. chenopodiifolia* reference transcriptome and also to the high-quality reference genome of the related diploid species *C. hirsuta* (Gan *et al*., 2016) (Fig. S3A). By combining these two approaches, we aimed to get the most out of our data set. On the one hand, the *C. chenopodiifolia* reference transcriptome provides perfect alignment, but it doesn’t represent the entire genome. On the other hand, alignment to *C. hirsuta* provides a complete diploid genome, but the alignment is not perfect.

Aligning the short read sequences from *C. chenopodiifolia* valves to the reference transcriptome of *C. chenopodiifolia* gave 46411 genes with a Uniprot ID after low count filtering, among which 31944 genes were unique (Table S1). By comparison, the *C. chenopodiifolia* short read sequences aligned to 16376 *C. hirsuta* genes, with 11870 genes in common to all four sample types, which corresponds to 40% of the annotated *C. hirsuta* genes (Fig. S3B). Only 18.6% of the short read sequences from *C. chenopodiifolia* fruit valves aligned to the *C. hirsuta* genome. Given that *C. chenopodiifolia* is octoploid, this fraction of the *C. chenopodiifolia* transcriptome that maps to *C. hirsuta* may suggest that *C. hirsuta* was a diploid progenitor species at some point during the evolutionary history of *C. chenopodiifolia*.

### Aerial and subterranean fruit show very different transcriptomes

Our experimental design allowed us to compare differentially expressed genes in both aerial and subterranean fruit valves between different stages of fruit development and between different fruit morphs (Fig. 5A). We reasoned that differentially regulated genes may be involved in processes such as endocarp *b* lignin patterning, collapse of the endocarp *a* layer, as well as photosynthesis, growth patterns of the fruit valves, or response to the contrasting environments of air versus soil. As such, this transcriptome analysis can provide a framework for future research on amphicarpy in *C. chenopodiifolia*.

By mapping *C. chenopodiifolia* short reads to the *C. hirsuta* genome, we identified differentially expressed genes (DEGs) based on an adjusted p-value of <0.05 and a minimum fold-change of 2, for the four possible comparisons: aerial fruit stage 16 (air16) vs aerial fruit stage 17ab (air17), 1752 DEGs; subterranean fruit stage 16 (sub16) vs subterranean fruit stage 17ab (sub17), 1076 DEGs; air16 vs sub16, 1939 DEGs; air17 vs sub17, 2859 DEGs, and 949 DEGs from the interaction of both fruit stages and fruit types (Fig. 5B-C, Tables S3-S5, Fig. S3C, H). We observed strong transcriptional changes for all comparisons, and in particular between the two fruit types at the later stage (17ab) (Fig. 5B-C). Indeed, most of the variance between samples can be explained by the difference in fruit types (Fig.5D, Fig. S4A). Many of the DEGs between fruit types were not differentially regulated across developmental stages (909 DEGs, Fig. 5B). These genes might represent distinct responses to aerial vs subterranean environments, or could also generally distinguish the two fruit types. A larger number of DEGs were affected by developmental stage in aerial than in subterranean fruit, and more DEGs were up-regulated than down-regulated in aerial compared to subterranean fruit (Fig. 5C). The same trends were found when mapping was performed with either the *C. hirsuta* genome or the *C. chenopodiifolia* reference transcriptome (Fig. S4B-G). Therefore, given that the annotation of the *C. hirsuta* genome is more complete, we used our alignment of short reads from *C. chenopodiifolia* fruit valves to the *C. hirsuta* genome for subsequent Gene Ontology (GO) analyses.

### Secondary cell wall formation is up-regulated and cell division is down-regulated during fruit valve development

One of the major events in both fruit types during the transition from stage 16 to 17ab is the deposition of a lignified secondary cell wall in the endocarp *b* cell layer of the valves. To assess whether our experimental design captured this, we performed GO analysis of common DEGs for these two comparisons (255 up- and 238 down-regulated DEGs in stage 16 compared to stage 17ab, irrespective of fruit morph, Fig.6A, Table S6). Enriched biological processes for up-regulated genes included secondary cell wall biogenesis, lignin/phenylpropanoid and xylan metabolic processes (Fig. 6B).

**Fig. 6.**
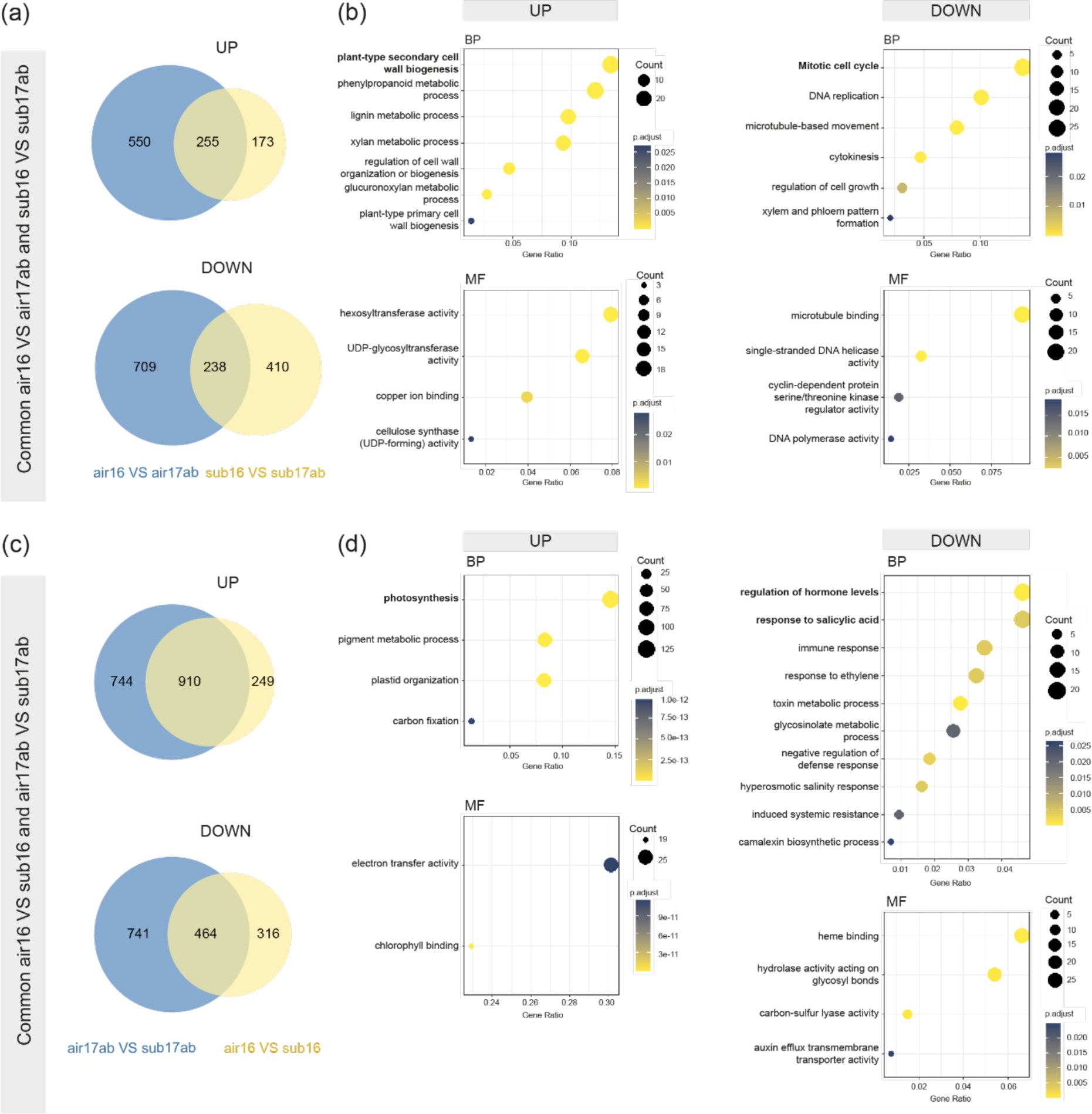
Gene Ontology (GO) enrichments for differentially expressed genes (DEG) in common to fruit development or fruit type. (a) Venn diagrams of significantly up- and down-regulated DEGs between stage 16 and stage 17ab aerial fruit compared to subterranean fruit (logFC ≥│1│, padj < 0.05). (b) Selected GO terms enriched in the 255 up-regulated DEGs and the 238 down-regulated DEGs that are in common to both fruit types, and therefore distinguish the two stages of fruit development. (c) Venn diagrams of significantly up- and down-regulated DEGs between aerial and subterranean fruit at stage 17ab compared to stage 16 (logFC ≥│1│, padj < 0.05). (d) Selected GO terms enriched in the 910 up-regulated DEGs and the 464 down-regulated DEGs that are in common to both developmental stages, and therefore distinguish the two fruit types. DEGs were obtained by mapping *C. chenopodiifolia* short reads to the *C. hirsuta* genome. Only genes with homologues in *A.thaliana* were included in the GO term analysis. BP, biological processes, MF, molecular function.

Molecular functions related to cell wall synthesis were also enriched, such as hexosyltransferase activity and cellulose synthase (UDP-forming) activity (Fig. 6B). Enrichment of copper ion binding GO terms could also indicate the involvement of copper-dependent laccase activity in endocarp *b* lignification, as reported for *C. hirsuta* (Fig. 6B) (Pérez Antón *et al*., 2022). This suggests that lignin and hemicellulose production for secondary cell wall formation dominate the processes occurring during these stages of valve development in both fruit types.

Both fruit types have already elongated by stage 17ab (Fig. 1H) and this is reflected in the enrichment of GO terms related to the cell cycle and cell growth in down-regulated genes (Fig. 6B). Other enriched GO terms such as DNA replication, DNA helicase activity, cytokinesis, microtubule-based movement and microtubule binding, support the idea that cell division and cell growth are downregulated by stage 17ab (Fig. 6B). Genes related to xylem and phloem pattern formation are also downregulated by stage 17ab (Fig. 6B), consistent with the development of vascular bundles in the fruit valve by this stage (Fig. 2B).

### Aerial fruit are characterized by photosynthesis, and subterranean fruit by hormone and immune responses

Growth in above-vs below-ground environments will inevitably affect the transcriptome of these two *C. chenopodiifolia* fruit morphs. To investigate this, we analysed common DEGs between aerial and subterranean fruit independent of developmental stage (910 up- and 464 down-regulated DEGs in aerial fruit compared to subterranean fruit, irrespective of stage, Fig. 6C). Photosynthesis was the most enriched biological process for genes up-regulated in aerial fruit, and other enriched processes and molecular functions were all related to photosynthesis (Fig. 6D, Table S6). Therefore, photosynthesis dominates the transcriptional landscape of green aerial fruit, compared to white subterranean fruit, due to their exposure to light.

Subterranean fruit were characterized by enriched GO terms for the regulation of hormone levels, including salicylic acid, ethylene, brassinosteroid and auxin, as well as for immune responses (Fig. 6D). Response to salicylic acid is an enriched term that may be associated with systemic acquired resistance (Fig. 6D) (Métraux *et al*., 1990). Induced systemic resistance is another enriched GO term, which acts via ethylene and jasmonic acid (Fig. 6D) (Pieterse *et al*., 2014). Response to ethylene is enriched, as are terms that are often associated with the jasmonic acid pathway, such as camalexin biosynthesis, glucosinolate and toxin metabolic processes (Fig. 6D). The abundance of microbiota in the soil may explain why immunity-related genes are up-regulated in subterranean vs aerial fruit. The enriched term hyperosmotic salinity response may also be explained by the soil medium being rich in different solutes (Fig. 6D). Taken together, the transcriptome of subterranean fruit is characterized by stress and hormonal responses, which may reflect their growth in a soil environment.

### A distinct set of cell wall-related genes are up-regulated during the development of aerial fruit valves

To identify candidate genes involved in developmental processes that distinguish the two fruit morphs, we analysed interactions between fruit stage and type (INTX, Fig. S3C). To this end, we clustered the sequence reads of DEGs according to their behaviours across samples (Fig. 7A) and performed a gene ontology analysis for each cluster (Fig. 7B, Table S7). We observed similar clustering using the sequence reads mapped to the *C. chenopodiifolia* reference transcriptome (Fig. S4H). The largest cluster contained genes with higher expression in aerial fruit (cluster #1, Fig. 7A). Genes related to photosynthesis were over-represented in this cluster and strongly down-regulated during the development of subterranean fruit valves (Fig. 7A-B). The fact that inhibition of photosynthetic gene expression was not just sustained but actually increased during development, suggests that the normal photosynthetic function of fruit valves (Brazel & Ó’Maoiléidigh, 2019) requires active repression in subterranean fruit (Fig.7B). The analysis of two additional clusters with similarly high expression in stage 17ab aerial fruit (clusters #1, #3 and #6, Fig. 7A), showed that all three clusters were enriched in genes involved in cell wall-related processes, including secondary cell wall and lignin biosynthesis (Fig.7B). Therefore, the up-regulation of cell wall-related genes during fruit development distinguishes aerial from subterranean fruit valves, suggesting that distinct genes may be associated with the different patterns of lignified secondary cell walls in these two fruit morphs.

**Fig. 7.**
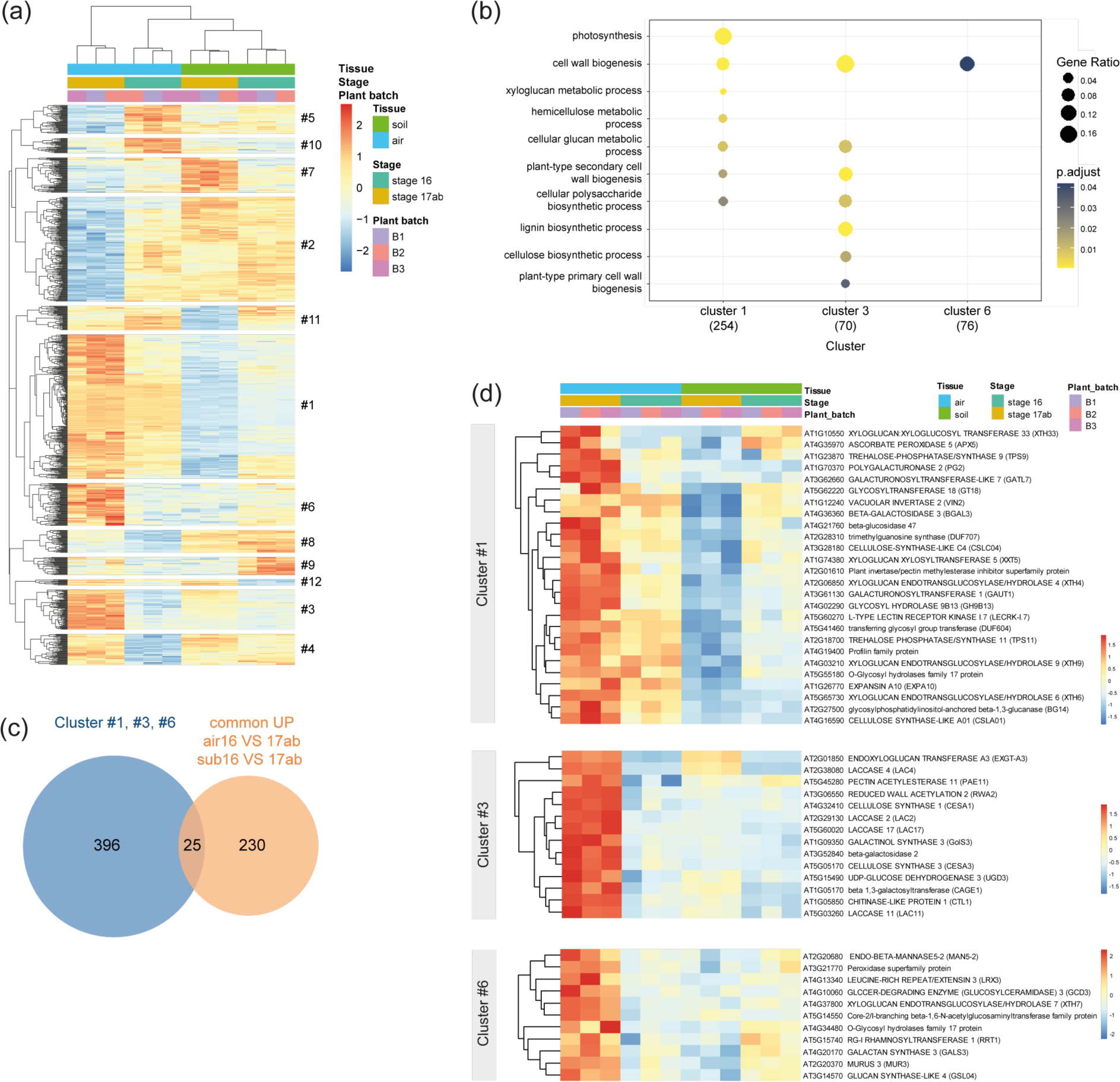
Differentially expressed genes (DEG) that distinguish the development of different fruit types. (a) Heat map of significant DEGs for the interaction comparison (INTX) between fruit stage and type (logFC ≥│1│, padj < 0.05). DEGs were obtained by mapping *C. chenopodiifolia* short reads to the *C. hirsuta* genome. (b) Selected gene ontology terms enriched in DEG clusters #1, #3 and #6 from (a). GO analysis was performed using *A. thaliana* GO term annotations for orthologous genes, in brackets. (c) Venn diagram of 421 DEGs in INTX clusters #1, #3, #6 from (b) compared with 255 common DEGs up-regulated in both aerial and subterranean fruit valves from Fig. 6a. (d) Heat maps of DEGs in INTX clusters #1, #3, #6 filtered for cell wall-related genes.

Since cell wall-related GO terms were also enriched in the set of DEGs common to the development of both fruit types (Fig.6A), we wondered if the same genes were identified in the interaction analysis (Fig. 7B). We found that only 25 genes overlapped between the sum of the three cell-wall related clusters (INTX #1, #3 and #6: 421 genes total) and the common genes up-regulated in both aerial and subterranean fruit valves (255 genes total) (Fig.7C). This indicates that the large majority of DEGs in the three cell-wall related clusters (INTX #1, #3 and #6) are specific to the development of aerial fruit. The 25 overlapping genes were all part of cluster #3 and characterized by enriched GO terms for lignin biosynthetic and metabolic processes, and cell wall biogenesis (Table S8). LACCASE 4 (LAC4/IRX12), LAC11 and LAC17, which are required for *C. hirsuta* endocarp *b* lignification (Pérez Antón *et al*., 2022), were included among these genes (Table S8). Therefore, these laccase genes are differentially expressed during the development of both explosive and non-explosive fruit, although to a much greater extent in explosive fruit.

To investigate the cell wall-related genes that are specific to aerial fruit, we filtered DEGs from each of the three INTX clusters (#1, #3 and #6) based on a list of genes annotated in *A. thaliana* that are related to primary cell wall biogenesis (Hartasánchez *et al*., 2023) (Fig.7D). Clusters #1 and #6 contained cell wall genes that are induced during aerial fruit development, but mostly repressed during the development of subterranean fruit (Fig. 7D). Among them, several genes are involved in hemicellulose synthesis or cell wall remodelling, such as XYLOGLUCAN ENDOTRANSGLUCOSYLASE/HYDROLASE (XTH) 4, 6, 7 and 9 (Fig. 7D). Cluster #3 contains some genes that are upregulated during the development of both fruit types (LAC2, LAC17, CAGE1 and ENDOXYLOGLUCAN TRANSFERASE A3), but all genes in this cluster are characterized by high expression in mature aerial fruit (Fig. 7D). These genes include CELLULOSE SYNTHASE (CESA) 1 and 3, which are part of the primary wall cellulose synthase complexes (Desprez *et al*., 2007; Persson *et al*., 2007). Whether up-regulation of cellulose or hemicellulose synthesis in the primary cell wall plays any role in patterning the distinctive endocarp *b* cell wall in explosive fruit are interesting questions for future research.

## Discussion

Plant trait diversity is an important resource. Focusing on plants that have not been well studied opens up the possibility to understand the development and evolution of different traits and the diversity of mechanisms by which plants successfully adapt to different environments. Amphicarpy is an under-studied trait that we investigate here in *C. chenopodiifolia.* This plant produces two types of fruit – above and below ground – with different seed dispersal strategies. We show that aerial, explosive fruit differ from subterranean, non-explosive fruit in morphology, lignin patterning and gene expression profiles related to the capacity for photosynthesis, secondary cell wall formation and defence responses. The tools generated here, together with a draft genome assembly (Emonet *et al*., 2024), establish *C. chenopodiifolia* as an emerging experimental system to study many distinctive developmental processes associated with amphicarpy.

Long read sequencing technologies enable accurate, high throughput characterization of the transcriptomes of under-studied species. Here, we took a hybrid approach, using PacBio IsoSeq long reads to define the isoforms expressed in the samples and short reads to get enough counts for well-powered differential expression. This cost-effective approach allowed us to compare the fruit valve transcriptomes of *C. chenopodiifolia* over development and between fruit morphs. By including IsoSeq long reads from additional tissue types and hormone/elicitor treatments in our reference transcriptome, we annotated a total of 53242 genes, enabling the expression of homeologous genes and species-specific genes or isoforms to be analysed. In this study, we chose to take advantage of the high-quality annotated reference genome of *C. hirsuta* to map *C. chenopodiifolia* short reads and facilitate DEG and GO analyses by enriching the annotation of gene functions. This approach captured similar trends between fruit morphs to what we found using the *C. chenopodiifolia* reference transcriptome, even though less than 20% of the short read data mapped to *C. hirsuta.* This alignment suggests that *C. hirsuta* may have contributed to the origin of *C. chenopodiifolia* as a diploid progenitor and tends to support an allopolyploidy origin of the *C. chenopodiifolia* genome (Emonet *et al*., 2024). These results highlight the advantages of selecting a study species within the Brassicaceae where model species already exist to enable comparative genomic and functional studies within and between species.

Cell wall biogenesis is a critical process that is upregulated during valve development in both fruit morphs. DEGs associated with this process are likely to reflect the growth of fruit valves and the deposition of lignified secondary cell walls in the endocarp *b* that occurs during these stages. Though many of these DEGs are common to the development of fruit valves in both morphs, we identified three large clusters that are strongly upregulated in aerial fruit at late stage (Fig.7A-B). Amongst these DEGs we identified *LAC4, 11* and *17*. These three laccases co-localise with the asymmetric pattern of lignin in endocarp *b* cells of explosive fruit in *C. hirsuta*, and are required to polymerize lignin in this secondary cell wall (Pérez Antón et al., 2022). In *C. chenopodiifolia*, *LAC4, 11* and *17* are differentially expressed in both fruit morphs, but their expression level is much higher in explosive aerial fruit. This finding suggests that *LAC4, 11, 17* may have a conserved function in endocarp *b* lignification that is not specific to explosive fruit and could be investigated in the non-explosive fruit of *A. thaliana*. In this way, comparative transcriptomics in *C. chenopodiifolia* fruit morphs provides a starting point to understand the genetic basis and the evolution of explosive dispersal.

Amphicarpy is a derived trait in the octoploid *C. chenopodiifolia* and polyploidy is often linked to the derivation of novel traits. There are many different ways in which whole genome duplication could lead to evolutionary novelties. The increased genetic variation and the buffering effect of duplicated genes can provide adaptive potential, particularly under environmental stress (Van de Peer *et al*., 2021). In allopolyploids, the merger of different traits from progenitor species could also lead to novelty (Sun *et al*., 2020). New traits could also arise from alterations in gene expression, epigenetic remodeling, sub-, and/or neofunctionalization of gene duplicates, and the rewiring of gene regulatory networks following whole genome duplication (Van de Peer *et al*., 2017). Both the reference transcriptome and draft genome of *C. chenopodiifolia* are useful resources to study polyploidy and trait evolution. For example, it will be interesting to determine whether the expression profiles of homeologous genes tend to be similar or divergent over development and between fruit morphs, and to more generally link behaviour of the transcriptome with the sub-genomes in *C. chenopodiifolia*.

Amphicarpy is a rare trait that provides unique insights into the development of very distinct reproductive structures in the above-versus below-ground environment. Flowers produced by the main shoot grow towards gravity and are well adapted to the soil environment. The strong and surprisingly long pedicels can dig the small flowers far below the surface and show circumnutation movements, which are known to help roots penetrate and grow around obstacles in the soil (Movie 1) (Taylor *et al*., 2021). These pedicels also have Casparian strip-like lignin deposition and it will be interesting to understand whether this has a barrier function similar to the root. The underground flowers remain closed and are particularly reduced in size, as often seen in cleistogamous flowers. These characteristics might help to limit damage as the flowers progress through the soil. Such a harsh environment also influences the transcriptome of the fruit valves, as genes involved in immune responses and salt stress are found to be up-regulated in these tissues. Roots are known to display a different immune response than the shoot in order to accommodate their microbiome (Wyrsch *et al*., 2015; Poncini *et al*., 2017). Whether subterranean flowers and fruits are specifically adapted to the soil environment would be an interesting question to follow up.

We also described two types of flowering mode in *C. chenopodiifolia*: rosette flowering in the apical shoot, and inflorescence flowering in the axillary shoots. Plants such as *A. thaliana* and *C. hirsuta* transition to the reproductive phase through inflorescence flowering. By contrast, rosette flowering plants such as *Leavenworthia crassa* do not bolt, but instead the apical meristem produces flowers supported by long pedicels directly at the centre of the rosette (Liu *et al*., 2011). *C. chenopodiifolia* produces its subterranean flowers in a similar way, although the pedicels are positively gravitropic and grow rapidly into the soil. Fruit burial by elongating pedicels seems unique to *C. chenopodiifolia*. Most amphicarpic plants rather bury fruit produced from aerial flowers or produce fruit directly on underground organs (Zhang *et al*., 2020a). Transforming the *L. crassa TFL1* gene into *A. thaliana* induced partial rosette flowering, coupled to the development of longer pedicels (Liu *et al*., 2011). This suggests that the elongated pedicels of subterranean flowers and rosette flowering of the apical shoot meristem may be linked traits in *C. chenopodiifolia*.

Stable genetic transformation in *C. chenopodiifolia* will be an essential tool to further investigate the many distinctive developmental processes associated with amphicarpy. We have provided a first proof of principle that Agrobacterium-mediated transformation by floral dip is possible, which paves the way for CRISPR/Cas9 gene editing and a suite of other molecular genetic approaches commonly used in model organisms. In addition, the phylogenetic relatedness of *C. chenopodiifolia* to other model species in the Brassicaceae, such as *A. thaliana* and *C. hirsuta*, allows inter-species gene transfer experiments to be applied as an evo-devo approach. More specifically, the *DR5v2::NLS-tdTomato* transgenic line that we report here allows auxin-related processes, such as the gravitropic response of subterranean flowers, to be investigated in detail.

In summary, by establishing *C. chenopodiifolia* as an experimental system, we provide the means to investigate the under-studied trait of amphicarpy: how it functions, how it evolved, and how resilient this bet-hedging strategy for seed dispersal may be to future climate change.

## Acknowledgements

We thank Peter Huijser for time-lapse photography, Rainer Franzen for scanning electron microscopy, Dagmar van Dusschoten and Daniel Pflugfelder for MRI measurements, Gregor Huber and Andreas Fischbach for their technical support and Hugo Hofhuis for preliminary observations. This work was supported by Swiss National Science Foundation fellowship P500PB_203021 to A.E. R.K. acknowledges support from the Helmholtz Association for the Forschungszentrum Jülich GmbH. A.H. acknowledges support from a Max Planck Society core grant to the Department of Comparative Development and Genetics.

## Author contributions

Conceptualization, A.E. and A.H.; Investigation, A.E., M.P.A., U.N.; Bioinformatics, S.D, A.E.; Resources, B.H. (PacBio sequencing), R.K. (Magnetic Resonance Imaging); Writing, A.E. and A.H.; Funding Acquisition, A.E. and A.H.; Supervision, A.H.

