## Supplemental Figures and Methods for "Amphicarpic development in the emerging model organism *Cardamine chenopodiifolia*"

The following Supporting Information is available for this article:

**Fig. S1** Development and plant architecture of *C. chenopodiifolia.*

**Fig. S2** Aerial vs subterranean fruit and seeds in *C. chenopodiifolia*.

**Fig. S3** RNA-seq workflow and analysis using mapping of *C. chenopodiifolia* short reads to the *C. hirsuta* genome.

**Fig. S4** RNA-seq analysis using mapping of *C. chenopodiifolia* short reads to the *C. chenopodiifolia* reference transcriptome.

**Table S1** *C. chenopodiifolia* PacBio IsoSeq reference transcriptome statistics.

**Table S2** *C. chenopodiifolia* Illumina short read sequence statistics.

**Table S3** Summary of differentially expressed genes in *C. chenopodiifolia* fruit valves mapped to the *C. hirsuta* genome or *C. chenopodiifolia* reference transcriptome.

**Table S4** Differentially expressed gene list from *C. chenopodiifolia* fruit valves using short reads mapped to the *C. chenopodiifolia* reference transcriptome.

**Table S5** Differentially expressed gene list from *C. chenopodiifolia* fruit valves using short reads mapped to the *C. hirsuta* genome.

**Table S6** Gene ontology analysis of differentially expressed genes in *C. chenopodiifolia* fruit valves.

**Table S7** Gene ontology analysis of differentially expressed genes in the interaction between fruit stage and fruit type in *C. chenopodiifolia* fruit valves. (Note: cluster #11 was too small to give significant results.)

**Methods S1**

**Movie S1** Time-lapse photography of *C. chenopodiifolia* subterranean fruit growing in soil over 26 days.

**Movie S2** Magnetic resonance imaging of *C. chenopodiifolia* subterranean fruit growth in soil.

**Movie S3** High speed movie of an exploding aerial seed pod in *C. chenopodiifolia* recorded at 10000 frames per second.

**Fig. S1 Development and plant architecture of *C. chenopodiifolia*.** (a) Plants at 5, 6, 7, 8.5, 10.5 and 12 weeks after germination. (b) Bar plot showing appearance of primary shoot (n=34), axillary shoot bolting (n=26), axillary shoot flowering (n=25) and explosive seed release (n=20) in days post-germination (dpg). (c) Box plot of number of rosette leaves (n=39) and axillary shoots (n=39) per plant, cauline leaves (n=35) and secondary branches (n=35) per shoot. (d) Transverse sections of aerial and subterranean pedicels stained for cellulose (Cyan, calcofluor white) and lignin (RedHot LUT, Basic fuchsin). Arrows indicate Casparian strip-like lignin deposition in the innermost cortical cell layer surrounding the vasculature. Scale bars: 10 cm (a), 50 μm (d).

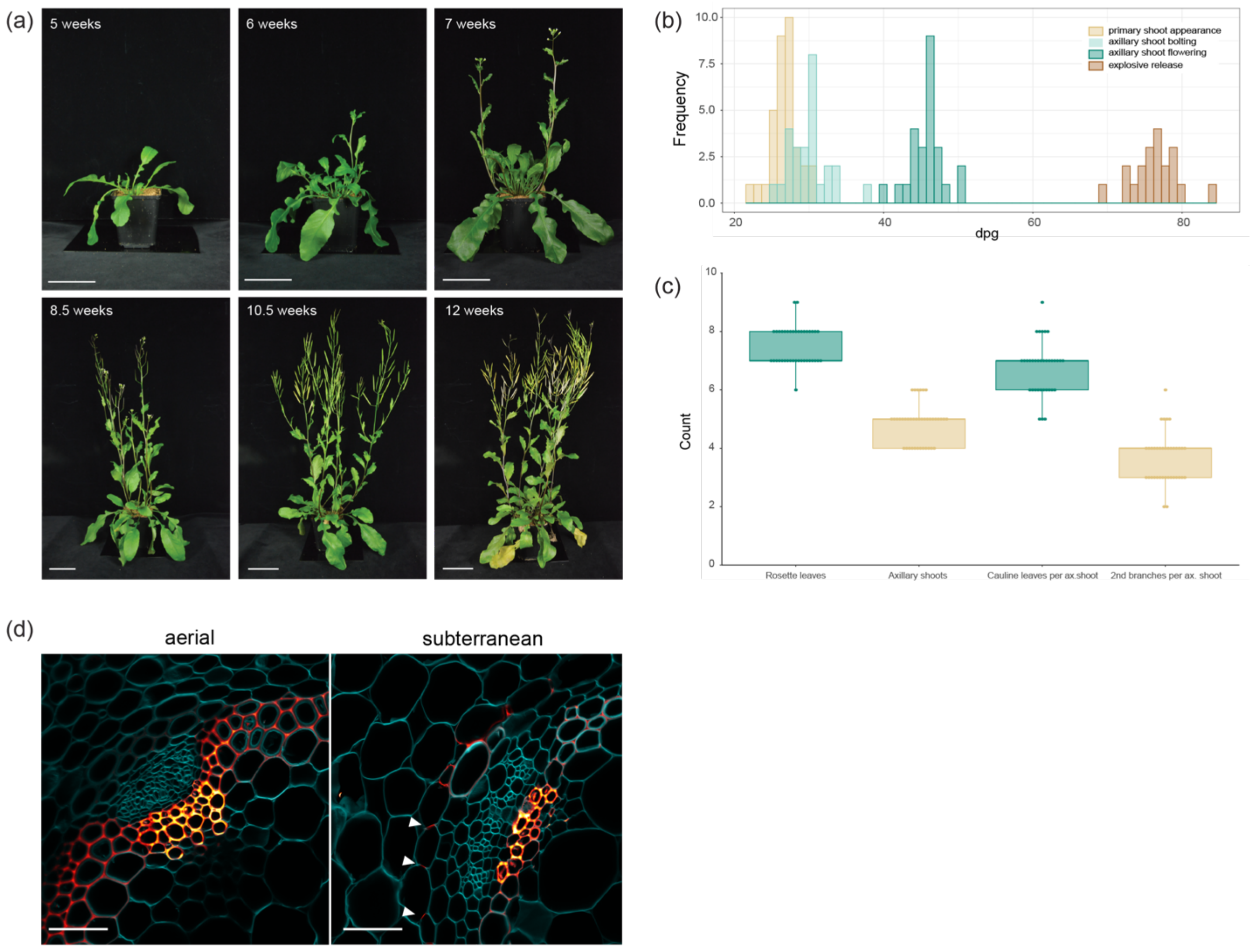

**Fig. S2 Aerial vs subterranean fruit and seeds.** (a-b) Valve margins (a) and replum (b) of aerial and subterranean fruit during stages 16, 17a, 17ab and 17b of fruit development. Transverse sections stained for cellulose (Cyan, calcofluor white) and lignin (RedHot LUT, Basic Fuchsin). (c-e) Scanning electron micrographs of aerial and subterranean seeds (c), seed surface (d) and funiculus (e). (f-g) Toluidine blue-stained transverse sections of (f) subterranean flower at stage 14 showing missing petals and fused stamens; S, sepals, P, petals; St, stamens; C, carpels; and (g) aerial and subterranean fruit at stage 17b showing seed coat, arrows indicate mucilage. Scale bars: 25 µm (a), 100 µm (b, e, f) 200 µm (c), 20 µm (b).

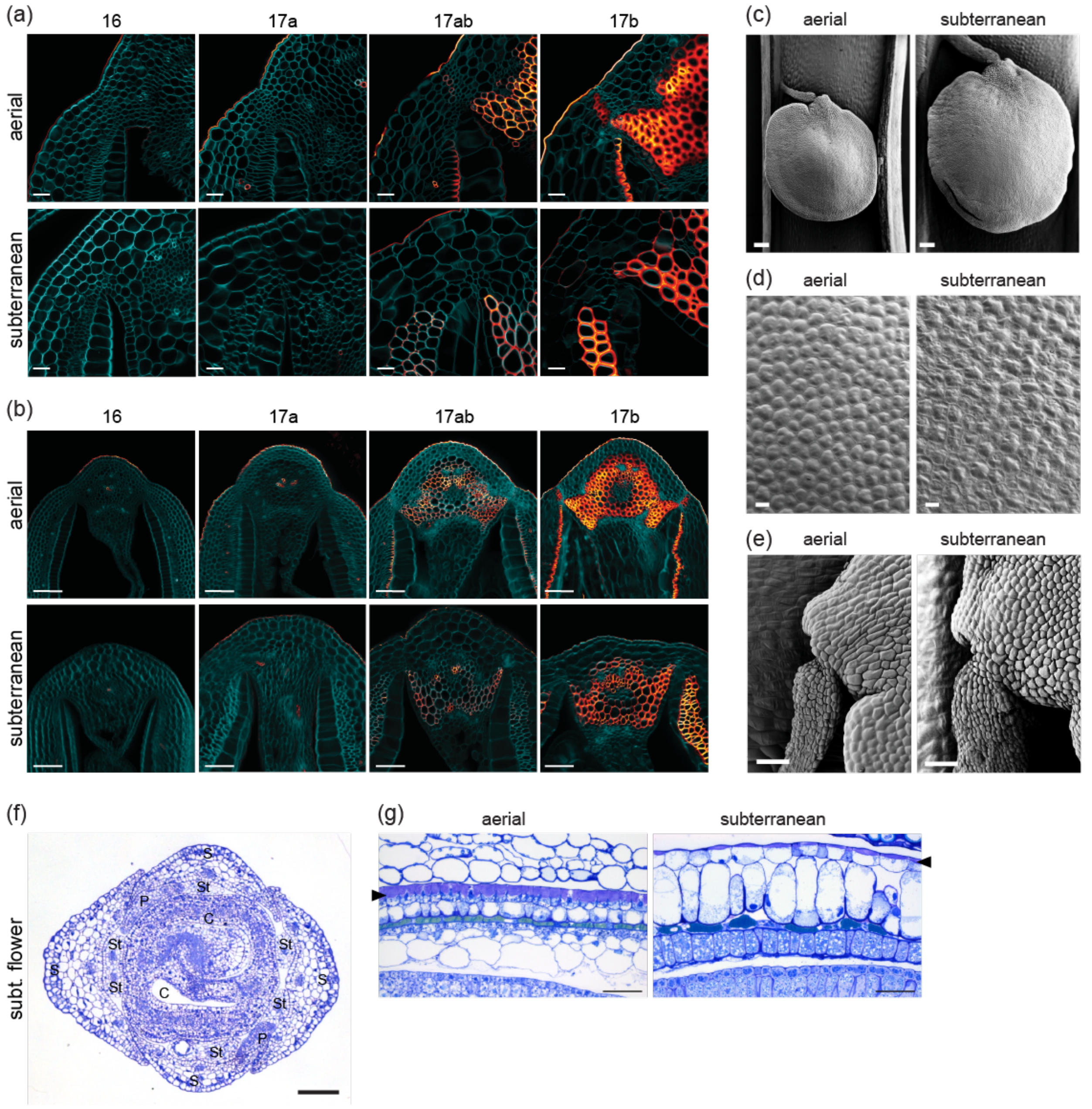

**Fig. S3 RNA-seq workflow and analysis using mapping of *C. chenopodiifolia* short reads to *C. hirsuta* genome.** (a) Analysis pipeline. (b) Venn diagram comparing genes identified in valves of aerial fruit stages 16 (air16) and 17ab (air17ab), and subterranean fruit stages 16 (sub16) and 17ab (sub17ab). (c-h) Venn diagram and volcano plots of DEGs ($logFC \geqslant |1|$, $padj < 0.05$) for all comparisons: air16 VS air17ab, air16 VS sub16, air17ab VS sub17ab, sub16 VS sub17ab, and the interaction between fruit stage and type (INTX). Blue dots, up-regulated DEGs ($logFC \geqslant 1$, $padj < 0.05$); red dots, down-regulated DEGs ($logFC \leqslant 1$, $padj < 0.05$).

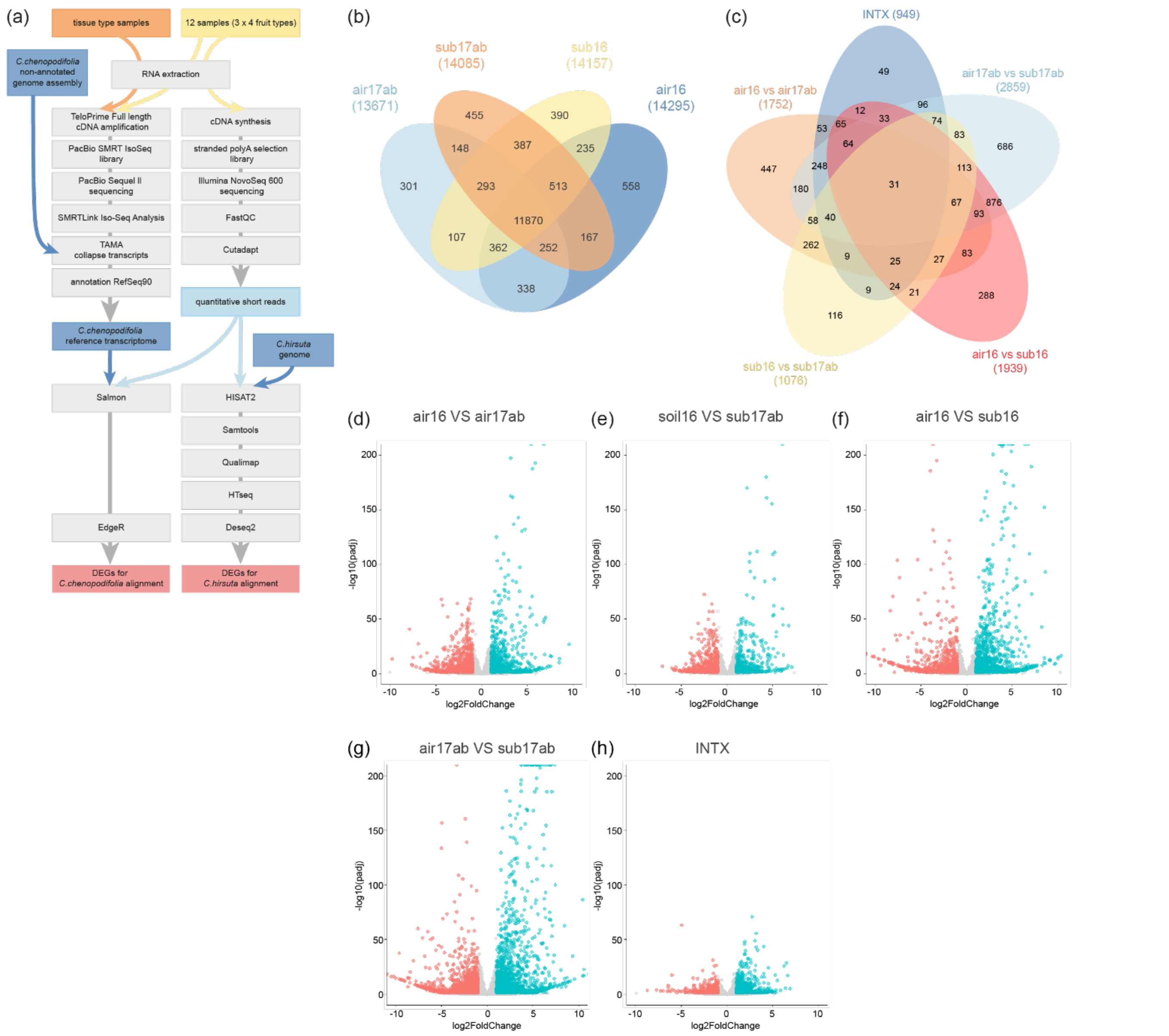

**Fig. S4 RNA-seq analysis using mapping of *C. chenopodiifolia* short reads to *C. chenopodiifolia* reference transcriptome.** (a) PCA of short read sequences from all four fruit valve sample types. (b-g) Venn diagram and volcano plots of DEGs (logFC ⩾ | 1 |, padj < 0.05) for all comparisons: air16 VS air17ab, air16 VS sub16, air17ab VS sub17ab, sub16 VS sub17ab, and the interaction between fruit stage and type (INTX). Blue dots, up-regulated DEGs; red dots, down-regulated DEGs. (h) Heat map of significant DEGs for the interaction comparison (INTX) between fruit stage and type.

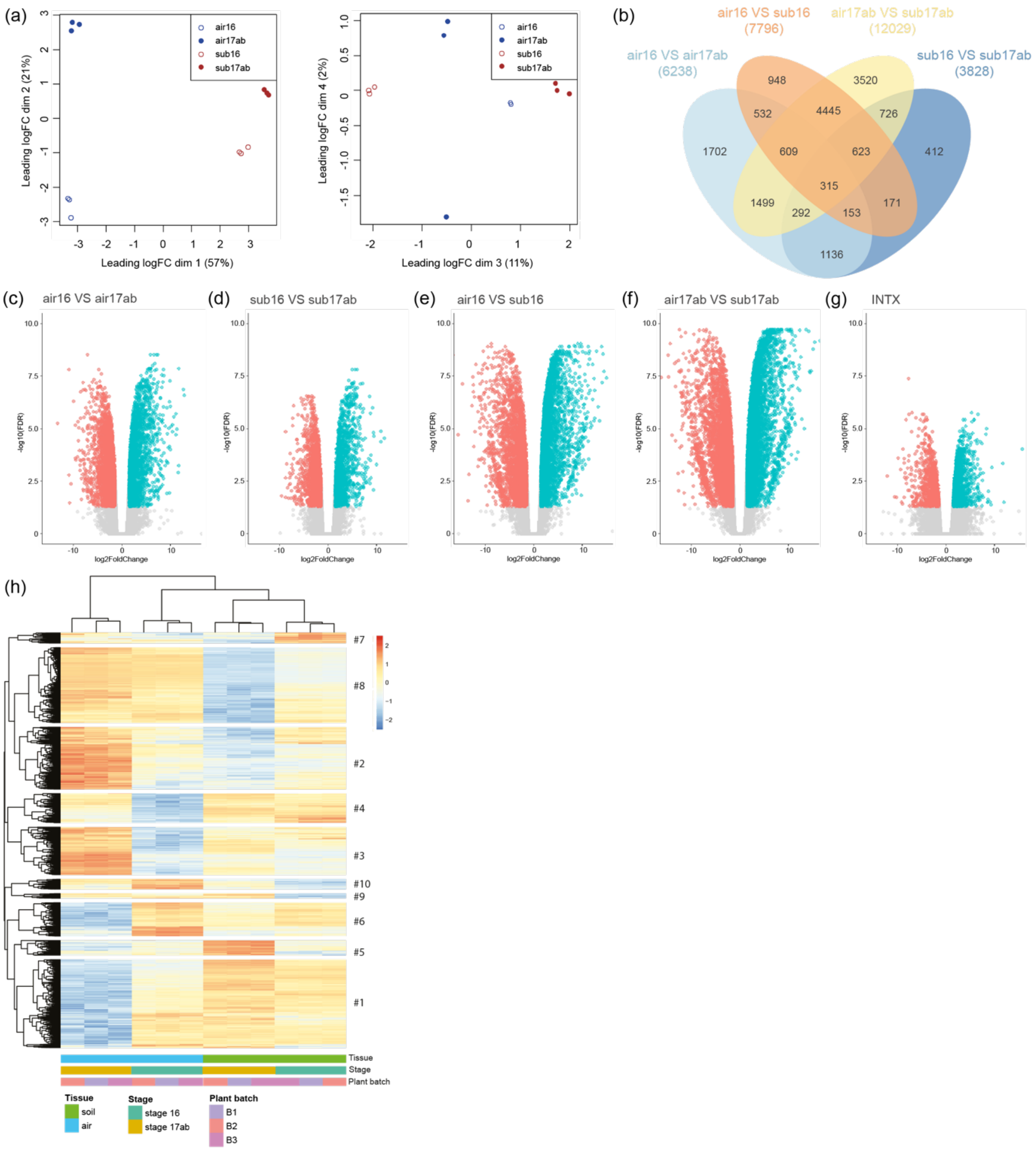

**Methods S1**

***Plant and growth conditions*** *Cardamine chenopodiifolia* aerial seeds were germinated *in vitro*. Seeds were sterilized in 70% EtOH with 0.1% Triton-X-100 for 20 min, rinsed in 100% EtOH and dried in an open Eppendorf under sterile conditions. Sterile seeds were sown on ½ Murashig and Skoog (MS), 1% agar (w/v) plates and stratified for 7 days in the dark at 4°C. Seedlings were grown in long day conditions (16h light, 20°C; 8h dark, 18°C) for seven days, then transferred to soil and grown in a walk-in growth chamber under long day conditions (16h light, 20°C; 8h dark, 18°C; 65% humidity). To assess flowering time under short day conditions, 1-week-old seedlings were transferred to a walk-in growth chamber set up with 8h light, 20°C; 16h dark, 18°C, 65% humidity. ***Plant transformation*** *DR5v2::NLS-tdTomato* construct in a modified pPZP200 vector was transformed in *Agrobacterium tumefaciens* GV3101 and a single colony inoculated in 5 ml LB supplemented with Spectinomycin (100 ug/ml) and Gentamycin (50 ug/ml) and shaken overnight at 28°C. One millilitre of this culture was inoculated into 1L LB supplemented with Spectinomycin and Gentamicin and shaken overnight at 28°C. At an OD of 0.8 to 1.0, the large culture was centrifuged in a Heraeus Multifuge 4KR Centrifuge for 20 min, the supernatant discarded and the pellet resuspended in 1L of a 50 g/L sucrose, ½ MS including vitamins, and 0.05% Silwet solution. Aerial flowers and stems of 7-week-old *C. chenopodiifolia* were immersed in this solution for around 1 min with agitation, and flower buds slightly manipulated to increase liquid penetration. Plants recovered overnight in the dark in a container covered with plastic foil before being transferred to the greenhouse. T1 seedlings were selected with basta treatment as follows: seeds were sown on wet soil and vernalized for 7 days at 4°C in the dark. Basta (0.2%) was sprayed every week for four weeks and resistant plants selected. ***Plant characterization and staging*** Forty plants were scored for: time after germination at first leaf appearance (n=30), time at first apical floral bud appearance (n=34), number of leaves at first apical floral bud appearance (n=32), time at axillary stem bud appearance (n=26), time at aerial flowering (n=25), total number of rosette leaves at time of bolting (n=39), total number of subterranean flowers and fruit (n=36), total number of axillary shoots (n=39), total cauline leaves on the first secondary branch (n=35), total secondary branches on the first axillary shoot (n=37), total number of aerial fruit on the first axillary shoot (excluding secondary branches) (n=30) and time at explosive shatter of first fruit (n=20). For staging, 17 plants at 44-53 dpg were dissected, aerial and subterranean fruit counted, their width and length measured and staged. Representative samples were sectioned and fixed

(Clearsee) for histology. ***Seed count, weight, and dimension analysis*** A MARVIN seed counter was used to count seeds and measure seed weight and dimensions of dry aerial and subterranean seeds from 4 plants. For aerial seeds, 3 batches of 100-200 seeds were measured per plant. Since subterranean seeds are less numerous, all of them were measured for each plant. To get the average number of seeds by fruit type, 10 stage 17b fruit/plant were dissected from 10 plants and the number of viable seeds counted. ***Mucilage assay*** To stain mucilage, seeds were imbibed with dd$H_2O$ for 2 hours at RT on an orbital shaker, stained with 0.01% ruthenium red solution for 1 hour, washed with dd$H_2O$ and observed under a Niko SMZ18 binocular with SHR Plan Apo 1.6x WD30 objective. ***High speed video*** Explosive pod shatter was filmed with a high-speed camera (Photron Fastcam SA4 120KM2) fitted with a 100 mm F2.8 lens. Aperture was set to f/4 and the software configured to save images at 10000 frames per second (fps). Fruit were illuminated with two DLED4.2-D Focusing LED Daylight Heads (DEDOTEC) and the camera activated manually after explosion was triggered using a wire mounted on electrical toothbrush or forceps to pinch the base of the fruit. ***Time lapse photography of subterranean fruit*** Seven-day-old *C. chenopodiifolia* seedlings were transferred to custom-made transparent chambers filled with soil (two 21x21cm Plexiglas sheets separated by 8 mm plastic spacer, mounted with steel clippers) and fixed in a tray with a capillary watering mat. The chamber was covered with aluminium foil to ensure dark conditions. Plants were grown for 4 weeks in long days until the first flower buds appeared, then the transparent chamber was unfoiled and inserted in a black plastic box designed to fit a Canon EOS 700D camera equipped with an EF-S 18-55mm f/3.5-5.6 lens, 2 flashlights with custom diffusers and an additional battery (PROPAC dual-output power source for flash PB960). The complete setup was transferred to a dedicated growth chamber (Hettich PRC1200SL growth cabinet illuminated with white-light (40%) plus FarRed light: 40%; PAR*: 160 µmol/sqm/s) and plants were grown for 26 days (growth conditions 16h/8h light/dark, 20°C/18°C; 60%/70% RH). Pictures were taken every 30 min with an exposure of 1/60s with ISO: 100 (total 1242 images). Images were processed as follows: RAW files were converted to TIF with Photoshop Camera Raw. FIJI function "Register Virtual Stack Slices" (Rigid -- translation + rotation) was then used before Sharpening and Rotation in Photoshop. Images were cropped to 5100x3400px. Rotation (clockwise) and size reduction to 75% were then applied, as well as crop to 2400x1600px. Final movie was obtained using FIJI after size reduction to 13.3px/mm (or 75µm/px), NanoJ-Core drift correction, Bleach correction – Histogram Match and final crop size AVI (1440x1080px; 1242 frames; 30.187min interval; 21.2358px/mm). The file was then reduced to 625 frames (averaging, 1 frame/hour), labelled, and exported as sequential frames. Conversion to .mp4 movie was done using Photoshop. ***MRI*** Two-week-old seedlings were transplanted to custom-made containers designed for MRI imaging (PVC tubes of 8 cm diameter and 14 cm height) with commercial field soil (loamy sand, sieved to 2 mm, brand name Sp2.1, Landwirtschaftliche Untersuchungs- und Forschungsanstalt Speyer, Speyer, Germany, http://tinyurl.com/m48bzlr) (Pflugfelder *et al.*, 2022). Plants were grown for 10 weeks in the greenhouse at MPIPZ until several below-ground fruits were produced. During cultivation, soil humidity was maintained constant at 70% of maximum water holding capacity (WHCmax). Magnetic resonance imaging (MRI) was performed at Jülich Forschungszentrum, Jülich, Germany. ***Histochemistry*** *Toluidine blue staining of resin-embedded sections:* 3mm segments of aerial and subterranean fruit at stages 17a and 17b were dissected in fixative medium while shoot apices and subterranean flowers were kept intact. Samples were fixed in 2.5% (v/v) glutaraldehyde (Agar Scientific, AGR1012), 2% paraformaldehyde (EM grade) in 0.05 M sodium cacodylate buffer (Agar Scientific, AGR1012, pH 6.9, diluted with MilliQ ELIX water) for 2 hours rotating at room temperature, dehydrated stepwise in ethanol over 2 days, and infiltrated stepwise with LR White over 48h. Samples were embedded in pure LR White in flat embedding molds at 60°C for 48 hours and 1.5 µm thin sections were stained with 0.05% toluidine blue. *Basic Fuchsin and Calcofluor white staining:* Fruit were embedded in 5% low melting agarose in a 1.5ml Eppendorf tube then 100-150µm transverse sections were cut using a Leica Vibratome VT1000 S. Sections were immediately fixed for 1-2 hours at room temperature in 4% paraformaldehyde in PBS solution, washed twice for 1 min with PBS, and cleared in Clearsee solution for at least 24 h with mild shaking. Sections were stained with 0.1% Calcofluor White and 0.2% Basic Fuchsin in Clearsee solution and incubated overnight. The staining solution was removed and samples rinsed once in fresh Clearsee solution, successively washed for 30 min and 2 hours in fresh Clearsee solution with gentle shaking before mounting in Clearsee for imaging. ***Microscopy*** *Binocular:* Nikon SMZ18 with ikon SHR Plan Apo 1.6x and 0.5x was used to image seed mucilage release. *Light microscopy:* Toluidine blue-stained sections were imaged using a Zeiss Axio Imager M2 upright epifluorescent microscope with EC Plan-Neofluar 10x/0.3, 20x/0.5 or 40x/0.75 objectives. *Confocal laser scanning microscopy:* Imaging was performed on a Leica TCS SP8 confocal scanning microscope. Pictures were taken with the HC PL FLUOTAR (10x/0.30 dry) and HCX PL APO lambda blue (63x/1.20 water) objectives. Excitation and detection windows were set as follows: for visualization of lignin and cellulose: calcofluor white (405nm, 425-450nm), basic fuchsin (561nm, 600-650nm). Images were processed using the Fiji software (Schindelin *et al.*, 2012). *Scanning electron microscopy:* Plant material was fixed overnight at 4°C in 4% glutaraldehyde in PBS (pH 7.2). Samples were washed twice in PBS, dehydrated stepwise in ethanol, and critical point dried using a Leica CPD300 critical point dryer. Samples were mounted, coated with platinum using a Polaron SC 7640 sputter coater, and imaged using a Zeiss Supra 40VP scanning electron microscope. ***RNA sequencing*** *Plant material and RNA extraction:* Aerial and subterranean fruit valves were collected from 7-week-old plants before lignification at stage 16-17a (aerial fruit: 1 mm width, 25 mm length; subterranean fruit: 2 mm width, 7 mm length) and during lignification at stage 17ab (aerial fruit: 2 mm width, 35 mm length; subterranean fruit: 3-4mm width, 12 mm length). For each fruit, the two valves were carefully dissected, taking care that no seeds, replum or septum tissue was included, and immediately flash-frozen in liquid nitrogen. For each stage 16 sample, 20 valves collected from 5 different plants (2 fruit per plant) were pooled together to ensure sufficient tissue. For stage 17ab samples, 10 valves (from 5 plants) were sufficient. The same 5 plants were used to collect the 4 different sample types. The experiment was repeated in triplicate. For the reference transcriptome, RNA was also collected from the following tissues: 5-day-old roots, 2-week-old shoot apices, 4-week-old flowering shoot apices, 4-week-old hypocotyl, 6-week-old flowering axillary shoot apices, aerial and subterranean fruit at stage 17ab, aerial and subterranean seeds. In addition, RNA was collected from 7-day-old seedlings treated for 2 h with 5 uM IAA, BA or flg22. Frozen tissues were ground using a TissueLyzer with 3 metal beads per tube, then total RNA extracted with Spectrum ™ Plant Total RNA kit (Sigma-Aldrich) following manufacturer's instruction. An additional DNase I digestion step was included and RNA was eluted in nuclease-free dd$H_2O$ before further processing. DNA with quantified by spectrophotometry (Nanodrop, Thermo Scientific, USA) and analysed by capillary electrophoresis using the Agilent 2100 Bioanalyzer (RNA Nanochip, Agilent Technologies, Germany). *Full length cDNA library preparation and PacBio SMRT long-read sequencing:* Full length cDNA synthesis was performed with the TeloPrime Full-Length cDNA Amplification Kit (Lexogen, Austria) (Cartolano *et al.*, 2016) and barcoded at the 5' end of the forward primer for multiplexing. Two single Molecule Real Time (SMRT) bell libraries were made as recommended by Pacific Biosciences (Palo Alto, U.S.A) with the Iso-Seq protocol, and the SMRT cells sequenced on PacBio Sequel II. Raw sequence data are available in ENA bioproject PRJEB69676. *cDNA library preparation and short-read Illumina sequencing:* cDNA synthesis, Stranded PolyA selection library preparation and two-sided, 150bp-sequencing were carried out by the Novogene company using the Illumina NovaSeq 6000 platform. Raw sequence data are available in ENA bioproject PRJEB69676. *Reference transcriptome and transcript function annotation:* SMRTLink (v10.2.0.133434) "Iso-Seq Analysis" was applied to demultiplex the PacBio Isoseq data and to obtain high-quality transcripts. Iso-Seq reads were used to build a *C. chenopodiifolia* reference transcriptome. High quality transcripts were collapsed using TAMA (tc_version_date_2019_11_19), *C. chenopodiifolia* genome assembly information and Minimap2 (Li, 2018) alignment program. *C. chenopodiifolia* genome assembly was obtained from https://www.ebi.ac.uk/ena/browser/view/PRJEB71776 (Emonet *et al.*, 2024). Annotation was performed with TAMA by predicting ORFs and comparing them to the Uniref90 protein databases. *Analysis of differentially expressed genes (DEG):* The *C. chenopodiifolia* reference transcriptome was used with Salmon v1.8.0 to estimate transcript abundance from Illumina reads (Patro *et al.*, 2017). The abundance estimates were then imported and transformed to gene level counts using tximport software (Soneson *et al.*, 2016). DEG analysis was performed using edgeR (Robinson *et al.*, 2010) to test for differences between the 4 different groups (sub_16, sub_17ab, air_16 and air_17ab), accounting for batch effects of extraction and plant batches and using as design formula: Design <- model.matrix (~0+group+Plants+Extraction). Contrasts between sample groups to test for differential expression were done for five comparisons: stage 16 VS stage 17ab in aerial fruit and subterranean fruit, respectively; aerial fruit VS subterranean fruit at stage 16 and 17ab, respectively; and the interaction of sample types and stages. The general linear model Quasi-likelihood F-tests (glmQLFTreat) was then used to find significantly DEGs between the groups. DEGs were selected based on a cut-off with a fold-change (FC) of 2, and a false-discovery rate (FDR) of 0.05. Alignment of Illumina reads to the *C. hirsuta* genome was done as follows. Short reads were quality controlled with FastQC and trimmed using the Cutadapt (Martin, 2011) software with default values, then aligned to the *C. hirsuta* genome (Gan *et al.*, 2016) using HISAT2 2.1.0. Samtools and Qualimap were used for conversion of BAM files and quality control before reads count analysis with HTSeq (García-Alcalde *et al.*, 2012; Danecek *et al.*, 2021). DEG analysis was carried out in R studio using DESeq2 package (Love *et al.*, 2014). Design was defined by the formula: design = ~ Tissue + Stage + Tissue:Stage. DEGs were selected based on FC >2 and adjusted p-value < 0.05. *Cluster and Gene Ontology analysis:* GO analysis was performed using *A. thaliana* GO term annotations for orthologous *C. hirsuta* genes (Gan *et al.*, 2016). Analyses were carried out in R using the package ClusterProfiler for gene ontology analysis (Wu *et al.*, 2021), ggplot2 for graphs and volcano plots generation (Wickham, 2016), pheatmap for heat maps (Kolde, 2019) and VennDiagram for Venn diagrams of DEG and reads counts (Chen & Boutros, 2011). Counts were normalized with the variance stabilizing transformation (vsd) methods for alignment to *C. hirsuta*. Venn diagrams were performed for all identified DEGs, while GO term analysis was performed for DEGs with homologues in Arabidopsis. ***Statistical analysis*** All statistical analyses were carried out using R (v4.2.1) and RStudio (v. 2022.07.1). Binary comparisons were performed using Student t-test. When the data didn't follow the linear model assumption, Wilcoxon Mann-Whitney test was used for binary comparison.

**Suplementary reference list**


**Cartolano M, Huettel B, Hartwig B, Reinhardt R, Schneeberger K**. **2016**. cDNA Library Enrichment of Full Length Transcripts for SMRT Long Read Sequencing. *PLOS ONE* **11**: e0157779.

**Chen H, Boutros PC**. **2011**. VennDiagram: a package for the generation of highly-customizable Venn and Euler diagrams in R. *BMC Bioinformatics* **12**: 35.

**Danecek P, Bonfield JK, Liddle J, Marshall J, Ohan V, Pollard MO, Whitwham A, Keane T, McCarthy SA, Davies RM, *et al.* 2021**. Twelve years of SAMtools and BCFtools. *GigaScience* **10**: giab008.

**Emonet A, Awad M, Tikhomirov N, Vasilarou M, Perez-Anton M, Gan X, Novikova PY, Hay A**. **2024**. Polyploid genome assembly of *Cardamine chenopodiifolia*. *bioRxiv*. doi: 2024.01.24.576990.

**Gan X, Hay A, Kwantes M, Haberer G, Hallab A, Ioio RD, Hofhuis H, Pieper B, Cartolano M, Neumann U, *et al.* 2016**. The *Cardamine hirsuta* genome offers insight into the evolution of morphological diversity. *Nature Plants* **2**: 1–7.

**García-Alcalde F, Okonechnikov K, Carbonell J, Cruz LM, Götz S, Tarazona S, Dopazo J, Meyer TF, Conesa A**. **2012**. Qualimap: evaluating next-generation sequencing alignment data. *Bioinformatics* **28**: 2678–2679.

**Kolde R**. **2019**. Pheatmap: pretty heatmaps.

**Li H**. **2018**. Minimap2: pairwise alignment for nucleotide sequences. *Bioinformatics* **34**: 3094–3100.

**Love MI, Huber W, Anders S**. **2014**. Moderated estimation of fold change and dispersion for RNA-seq data with DESeq2. *Genome Biology* **15**: 550.

**Martin M**. **2011**. Cutadapt removes adapter sequences from high-throughput sequencing reads. *EMBnet. journal* **17**: 10–12.

**Patro R, Duggal G, Love MI, Irizarry RA, Kingsford C**. **2017**. Salmon provides fast and bias-aware quantification of transcript expression. *Nature Methods* **14**: 417–419.

**Pflugfelder D, Kochs J, Koller R, Jahnke S, Mohl C, Pariyar S, Fassbender H, Nagel KA, Watt M, van Dusschoten D**. **2022**. The root system architecture of wheat establishing in soil is associated with varying elongation rates of seminal roots: quantification using 4D magnetic resonance imaging. *Journal of Experimental Botany* **73**: 2050–2060.

**Robinson MD, McCarthy DJ, Smyth GK**. **2010**. edgeR: a Bioconductor package for differential expression analysis of digital gene expression data. *Bioinformatics* **26**: 139–140.

**Schindelin J, Arganda-Carreras I, Frise E, Kaynig V, Longair M, Pietzsch T, Preibisch S, Rueden C, Saalfeld S, Schmid B, *et al.* 2012**. Fiji: an open-source platform for biological-image analysis. *Nature Methods* **9**: 676–682.

**Soneson C, Love MI, Robinson MD**. **2016**. Differential analyses for RNA-seq: transcript-level estimates improve gene-level inferences. *F1000Research* **4**: 1521.

**Wickham H**. **2016**. Data Analysis. In: Wickham H, ed. Use R! ggplot2: Elegant Graphics for Data Analysis. Cham: Springer International Publishing, 189–201.

**Wu T, Hu E, Xu S, Chen M, Guo P, Dai Z, Feng T, Zhou L, Tang W, Zhan L, *et al.* 2021**. clusterProfiler 4.0: A universal enrichment tool for interpreting omics data. *The Innovation* **2**: 100141.
